# Reversible histone deacetylase activity catalyzes lysine acylation

**DOI:** 10.1101/2023.11.17.567549

**Authors:** Takeshi Tsusaka, Mohd. Altaf Najar, Benjamin Schwarz, Eric Bohrnsen, Juan A. Oses-Prieto, Christina Lee, Alma L. Burlingame, Catharine M. Bosio, George M. Burslem, Emily L. Goldberg

**Affiliations:** Department of Physiology, University of California, San Francisco; San Francisco, CA, 94158, USA; Department of Biochemistry and Biophysics, Perelman School of Medicine, University of Pennsylvania; Philadelphia, PA, 19104, USA; Immunity to Pulmonary Pathogens Section, Laboratory of Bacteriology, National Institute of Allergy and Infectious Diseases; Hamilton, MT, 59840, USA; Department of Pharmaceutical Chemistry, University of California, San Francisco; San Francisco, CA, 94158, USA; Department of Molecular and Cell Biology, University of California, Berkeley; Berkeley, CA, 94720, USA; Department of Cancer Biology and Epigenetics Institute, Perelman School of Medicine, University of Pennsylvania; Philadelphia, PA, 19104, USA; Chan-Zuckerberg Biohub; San Francisco, CA, 94158, USA

## Abstract

Starvation and low carbohydrate diets lead to the accumulation of the ketone body, β-hydroxybutyrate (BHB), whose blood concentrations increase more than 10-fold into the millimolar range. In addition to providing a carbon source, BHB accumulation triggers lysine β-hydroxybutyrylation (Kbhb) of proteins via unknown mechanisms. As with other lysine acylation events, Kbhb marks can be removed by histone deacetylases (HDACs). Here, we report that class I HDACs unexpectedly catalyze protein lysine modification with β-hydroxybutyrate (BHB). Mutational analyses of the HDAC2 active site reveal a shared reliance on key amino acids for classical deacetylation and non-canonical HDAC-catalyzed β-hydroxybutyrylation. Also consistent with reverse HDAC activity, Kbhb formation is driven by mass action and substrate availability. This reverse HDAC activity is not limited to BHB but also extends to multiple short-chain fatty acids. The reversible activity of class I HDACs described here represents a novel mechanism of PTM deposition relevant to metabolically-sensitive proteome modifications.

---

Glucose is a primary fuel source for energy production across species. When restricted during prolonged fasting or when consuming a low-carbohydrate diet, an adaptive starvation response is triggered to increase ketone body production by the liver (1). Ketone bodies are short-chain fatty acids (SCFA) that are utilized in tissues such as the brain, heart, and skeletal muscle to generate acetyl-coenzyme A (CoA) and for ATP production in a glucose-depleted state (2). Consequently, ketone concentrations are highly dynamic, increasing more than 10-fold to reach 1-3 mM in circulation during fasting.

The most abundant ketone body is β-hydroxybutyrate (BHB). In addition to its role in ATP generation, this ketone has been implicated in other cellular functions (3). Prior studies show that BHB actively inhibits histone deacetylases (HDACs) (4) to affect gene expression. Others show that BHB stimulates G protein-coupled receptors at physiological concentrations (5, 6). Most recently, BHB was identified as a novel protein modification on lysine, an adduct known as Kbhb (7). The abundance of Kbhb modifications across the proteome is proportional to BHB concentration both in cell culture and in vivo in mice (7, 8). Kbhb is found on numerous proteins in nearly every cellular compartment, and many histone Kbhb sites are residues that are targeted for other acylations (9). However, the machinery responsible for the deposition of this post-translational modification (PTM) remains poorly understood.

Lysine acylations can occur non-enzymatically via direct reaction with acyl-CoA, are “written” enzymatically by lysine acyltransferases, and are “erased” by lysine deacylases (10-15). Histone deacetylases (HDACs) are a conserved family of enzymes structurally divided into four classes, with Class I, II, and IV being Zn2+-dependent, and the Class III Sirtuins being NAD+-dependent (16). While acetylation is the most prevalent and well-studied lysine acylation, the basic principle of PTM control extends to other acyl groups, including lactate (17, 18) and β-hydroxybutyrate (9). Using biochemical and mass spectrometry approaches, we set out to identify additional regulators of Kbhb formation. As reported below, we discover a non-canonical role for class I HDACs (HDACs 1, 2, and 3) with the ability to reverse their activity and catalyze the formation of acylated lysine. Moreover, the same residues in the active site of class I HDAC are required for both deacetylation and the non-canonical, reversible reaction β-hydroxybutyrylation. These data identify a new function of this enzyme class and a new mechanism of PTM regulation.

## Kbhb formation requires class I HDACs

To gain insights into the regulation and potential significance of Kbhb, we treated HEK293T cells with BHB and immunoprecipitated modified proteins with a validated pan-Kbhb antibody (Fig. 1, fig. S1, table S1) for global LC-MS/MS analysis. We used SAINT (*19*) analysis to identify significantly enriched proteins within the Kbhb proteome and found HDAC1 and HDAC2 as well as known HDAC1/2-complex proteins including MTA1-3, RBBP4/7, SIN3A, and NCOR1 (Fig. 1A,B), which were recently reported to remove Kbhb marks (*9*). We verified Kbhb formation on HDAC2 in BHB-treated cells by immunoblotting for Kbhb after FLAG-IP in cells engineered to express 3xFLAG-mHDAC2 (Fig. 1C). To explore the relationship between class I HDACs and Kbhb in more detail, we treated cells with HDAC chemical inhibitors trichostatin A (TSA), suberoylanilide hydroxamic acid (SAHA, clinically known as Vorinostat), MS-275, and butyrate (*20*) and found that each of these also blocked Kbhb formation (Fig. 1D, fig. S2A). Importantly, we observed similar results in several different human and murine cell lines (fig. S2B,C). Given that the only HDACs targeted by all four inhibitors are class I HDACs 1, 2, and 3, we used siRNA to knockdown each of these proteins in HEK293T cells to test their individual contributions to Kbhb formation (Fig. 1E). Strikingly, Kbhb induction was almost completely abolished when all three HDACs were knocked down, with HDAC2 having the single-greatest effect on Kbhb formation. We observed similar reduction in Kbhb in cells lacking HDAC1 or HDAC2 using CRISPR/Cas9 (fig. S2D) although we were unable to generate HDAC1/2 double knockout cells. These data indicate that HDACs 1, 2, and 3 redundantly contribute to the formation of Kbhb through an unknown mechanism.

**Fig. 1.**
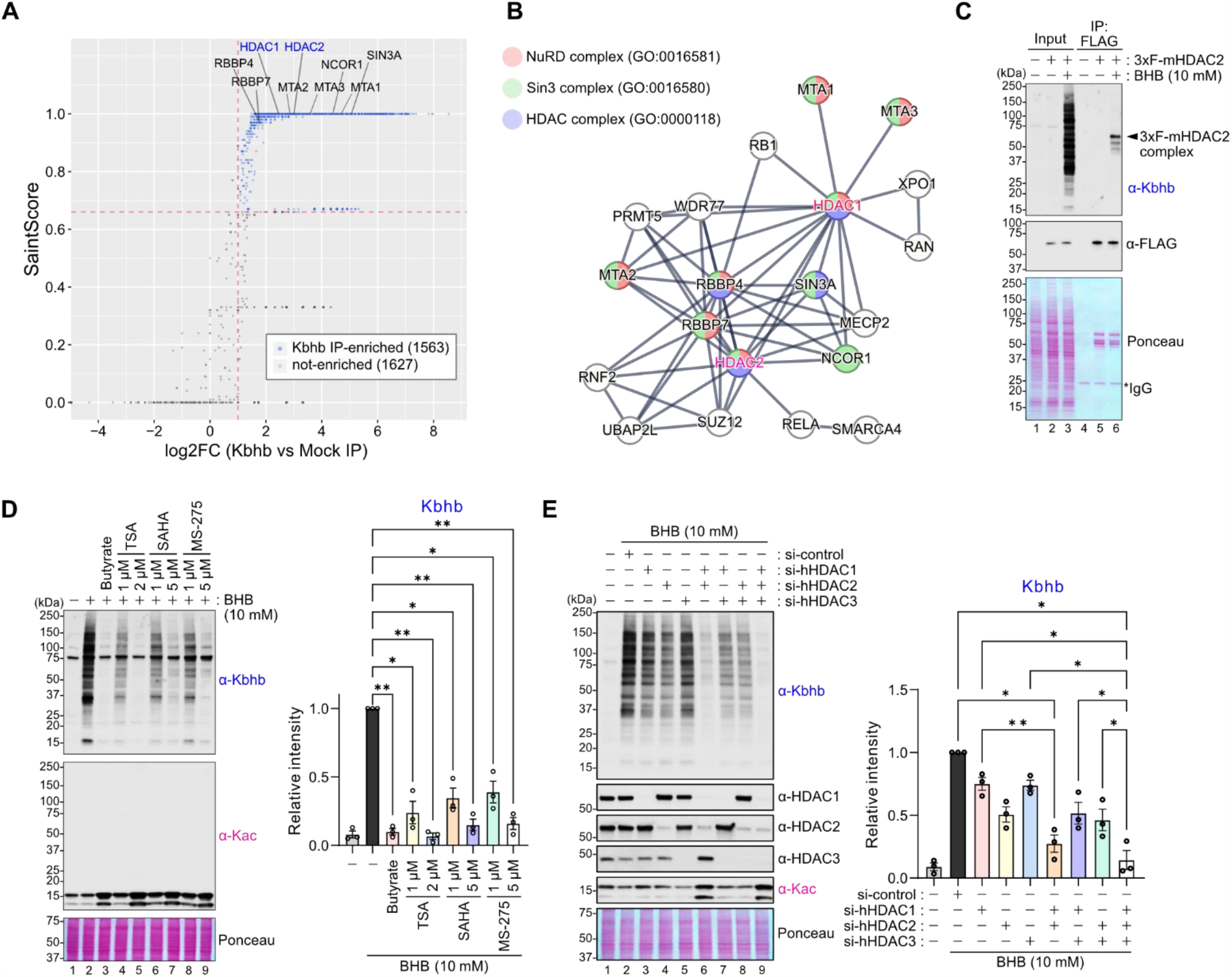
Class I HDACs are enriched in the Kbhb proteome and are required for Kbhb formation. (**A**) Scatter plot of Saint Score and log2(fold-change; FC, Kbhb vs Mock IP). Kbhb IP-enriched proteins were determined based on criteria as indicated in the Materials and Methods and are indicated in blue. Dots for HDAC-related proteins are indicated with their name. (**B**) The network of HDAC-related proteins in the Kbhb proteome are visualized by STRING analysis. Proteins that are in known HDAC complexes are highlighted in the corresponding color. (**C**) Validation of Kbhb modification on the HDAC2 complex. HDAC2 KO HEK293T cells expressing 3xFLAG-mHDAC2 was treated with or without 10 mM BHB for 24 hours and used for immunoprecipitation with anti-FLAG beads. Input and IPed samples were analyzed by western blot for Kbhb modification. (**D**) Left: Representative western blots of HEK293T cells treated with 10 mM BHB in combination with each HDAC inhibitor at the indicated concentrations for 24 hours. Representative of N=3 independent experiments. Right: Relative intensities of anti-Kbhb signals to the lane 2 (10 mM BHB treatment). Signals were normalized to ponceau S staining. Data are represented as mean +/-SEM of independently performed experiments and each symbol represents an individual experiment. Statistical differences were calculated by 1-way ANOVA followed by Dunnett’s correction for multiple comparisons. (**E**) Left: Representative western blots of HEK293T cells transfected with siRNA(s). Cells were treated with 10 mM BHB for 4 hours. Representative of N=3 indpendent experiments. Right: Relative intensities of anti-Kbhb signals to the lane 2 (10 mM BHB treatment). Signals were normalized by ponceau S staining. Data are represented as mean +/-SEM and each symbol represents an independently performed experiment. Statistical differences were calculated by 1-way ANOVA followed by Tukey’s test for multiple comparisons. *p<0.05, **p<0.01.

## Canonical lysine acylation pathways are not required for Kbhb formation

The requirement of HDACs for Kbhb modification prompted us to reevaluate the proposed mechanism of β-hydroxybutyrylation proceeding through a BHB-CoA-dependent pathway (fig. S3A) (*9*). Using cellular extracts, purified bovine serum albumin (BSA), or histone H3 as model substrates, we confirmed in vitro that BHB-CoA is sufficient for non-enzymatic formation of Kbhb, but this was not inhibited by butyrate (fig. S3B,C). Next, we ruled out the possibility that butyrate blocked BHB entry into cells with targeted metabolomics (fig. S3D,E). However, in these analyses we were unable to detect BHB-CoA even after high-dose BHB treatment (5 mM for 24 hours), and this was in contrast to butyryl-CoA, which was significantly elevated in butyrate-treated cells (fig. S3F,G). Moreover, siRNA-mediated knockdown and chemical inhibition of numerous acyl-CoA synthetase enzymes had no impact on Kbhb abundance in HEK293T cells (fig. S3H-L). These data do not support a role for BHB-CoA as a primary mechanism of Kbhb formation in cells.

Given that BHB-CoA may not be the intracellular BHB donor for Kbhb formation, we investigated whether acyltransferases might be involved in this acylation. In contrast to what was previously shown for specific histone Kbhb sites, we were unable to find a role for p300 or CBP in Kbhb formation broadly across the Kbhb proteome (fig. S4) (*9, 21*). Altogether, our data suggest an alternative mechanism of β-hydroxybutyrylation, independent of an acyl-CoA intermediate, exists and requires HDACs 1, 2, and 3.

## HDACs 1, 2, and 3 are sufficient to catalyze Kbhb formation

We first considered the possibility that global loss of HDAC activity might indirectly prevent Kbhb formation if, for instance, the target lysine residues consequently retained acetylation marks. However, we found no evidence of corresponding global Kac induction when we treated cells with HDAC inhibitors (Fig. 1D, fig. S2A-C). In addition, our Kbhb proteome analysis recovered hundreds of proteins across all cellular compartments from BHB-treated HEK293T cells (fig. S1), further decreasing any likelihood that Kbhb sites would be occupied by other PTMs.

Since many enzymes catalyze reversible processes, we next considered the hypothesis that HDACs might directly catalyze protein β-hydroxybutyrylation. To test this hypothesis, we developed an in vitro reconstitution assay with recombinant human HDAC2 (rHDAC2), BHB, and recombinant histone H3.1 (rH3) as a model bait protein (Fig. 2A). Western blot analysis showed that rH3 is β-hydroxybutyrylated only in the presence of both rHDAC2 and BHB. The β-hydroxybutyrylation by rHDAC2 is BHB dose-dependent (fig. S5A) and occurs at physiological pH 6.5-8.5 (fig. S5B). Time-course analysis shows that the β-hydroxybutyrylation on rH3 can occur within 5 minutes and plateaus by 60 minutes under these conditions (fig. S5C). Pre-heating rHDAC2 or adding the HDAC active site inhibitor TSA eliminates Kbhb formation, indicating enzymatic activity of HDAC2 is necessary for target protein β-hydroxybutyrylation (Fig. 2B,C). To verify that this reaction is not specific to H3, we performed the same experiment with 60°C-inactivated whole cell lysates and obtained similar results (fig. S5D). We also confirmed these data with rHDAC2 purchased from a different vendor (Fig. S6). In addition to HDAC2, we investigated whether other HDACs could similarly catalyze Kbhb formation. We tested commercially available rHDACs 1-8, except for rHDAC7, in our in vitro reconstitution assay (Fig. 2D,E). Consistent with our HDAC siRNA knockdown experiment (Fig. 1C) we found that only HDACs 1, 2, and 3 had β-hydroxybutyrylation capability. These HDACs also have stronger deacetylation activity than other HDACs (Fig. 2F), as previously reported (*22, 23*).

**Fig. 2.**
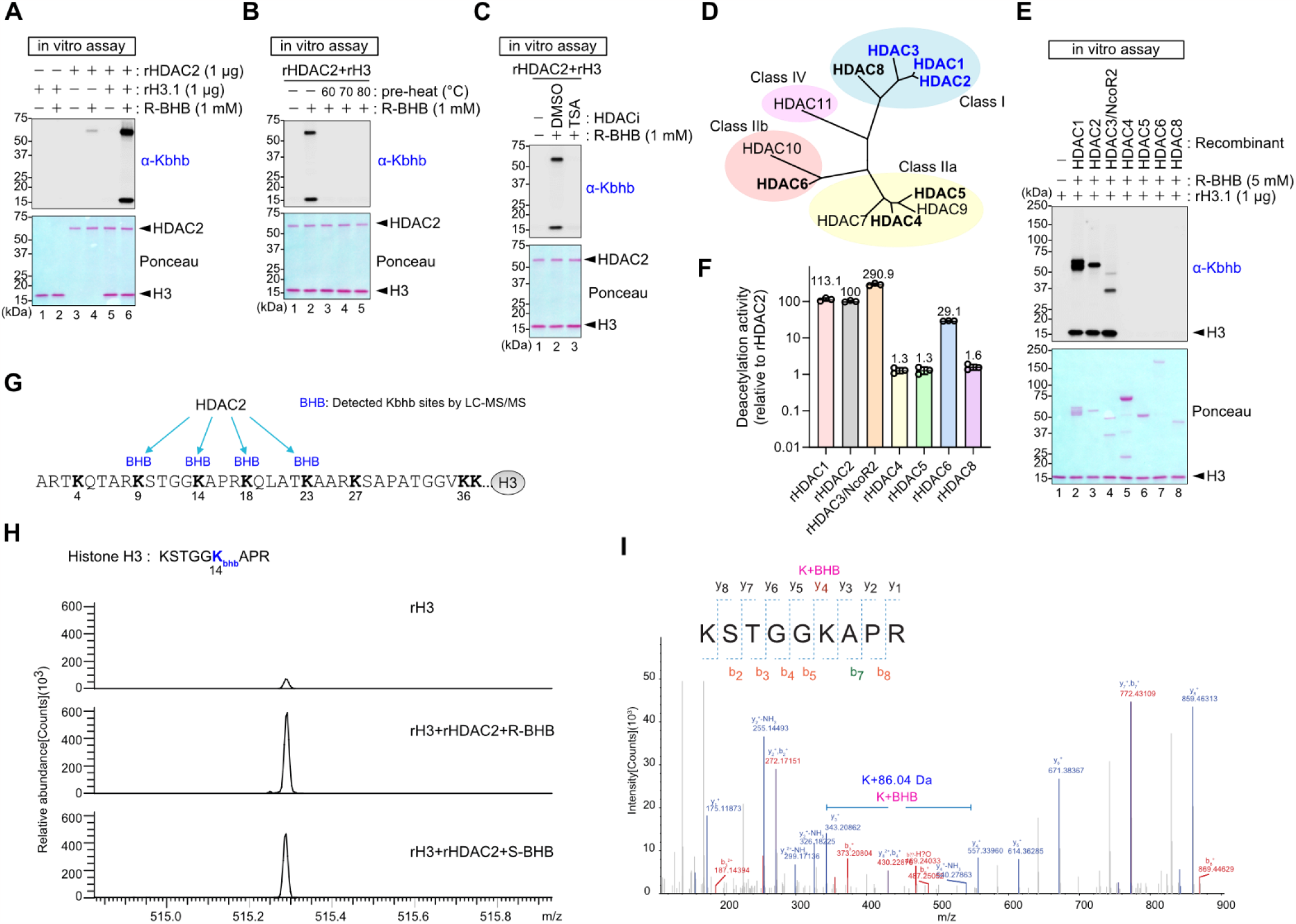
HDAC1, 2, and 3 catalyze lysine β-hydroxybutyrylation vitro. (**A-C**) Western blots of an in vitro lysine β-hydroxybutyrylation assay with recombinant HDAC2 (rHDAC2) and histone H3 (rH3) and R-BHB. Reaction was performed at 37°C for 1 hour. Protein loading was visualized by ponceau S staining. Kbhb was detected by anti-Kbhb antibody. rHDAC2 was inactivated by heating for 1 hour at the indicated temperatures in (B). TSA at 5 μM was added in the reaction mixture in (C). (**D**) Phylogenic tree of human HDAC1-11. Recombinant proteins used in (E) and (F) are indicated in bold. HDACs that are capable of lysine β-hydroxybutyrylation in (E) are highlighted in blue. (**E**) Representative western blots of an in vitro lysine β-hydroxybutyrylation assay with rHDAC1-8 (except for HDAC7) and rH3 and R-BHB. Approximately 1 μg HDACs were used for each reaction. (**F**) Relative deactylation activity of each rHDAC as compared to rHDAC2 was measured by fluorogenic peptide deacetylation assay. The same amounts of rHDACs as in (E) were used. (**G-I**) LC-MS/MS analysis on H3 after in vitro lysine β-hydroxybutyrylation assay. Similar results were obtained from N=2 independent experiments. Detected Kbhb modified sites on H3 are summarized in (G). MS-spectrum of the indicated peptide is shown in (H). Both R- and S-BHB induced Kbhb on lysine 14. MSMS-spectra for the indicated peptide with detected y and b ions are shown in (I).

Based on our previous study that some histone site-specific Kbhb antibodies are non-specific (*24*), we used liquid chromatography– tandem mass spectrometry (LC-MS/MS) to verify lysine β-hydroxybutyrylation on rHDAC2 and rH3. These analyses confirmed a chemical adduct of 86.04 Da on multiple lysine residues, as expected (Fig. 2G-I, fig. S7, table S2). As the pan-Kbhb antibody is R-BHB enantiomer-specific and cannot detect S-BHB (*7*), we used our purified reconstitution assay with LC-MS/MS and discovered that HDAC2 can catalyze β-hydroxybutyrylation with both R- and S-BHB enantiomers. Collectively, our data show that select HDACs are capable of β-hydroxybutyrylating target proteins.

## HDAC β-hydroxybutyrlation capability is coupled to deacetylation activity

We next sought to identify the molecular mechanism of β-hydroxybutyrylation by HDAC2 to determine if we could uncouple it from canonical deacetylation activity. Based on prior studies that identified several key residues for deacetylation (*25*), we generated HDAC2 KO HEK293T cell lines complemented with stable expression of 3xFLAG-tagged wild type (WT) or single-point mutants of murine HDAC2 (mHDAC2) (Fig. 3A). Specifically, we targeted residues involved in zinc ion binding or charge-relay, including H141A, H179A, D265N, and Y304F, and residues in the acetate escape channel, including R35K and C152A (*26*). When we expressed these proteins in HEK293T cells, combined with siRNA to knockdown HDACs 1 and 3 (to further enhance HDAC2-dependent Kbhb signal-to-noise), cells complemented with mutant mHDAC2 had lower Kbhb abundance and correspondingly higher Kac in whole-cell lysates (fig. S8). To evaluate the importance of each mutant more directly, we immunoprecipitated FLAG-tagged mHDAC2 to use in our Kbhb in vitro reconstitution assay (Fig. 3B). We found that WT 3xFLAG-mHDAC2 induced the most Kbhb on H3 and all mutants showed different degrees of lower β-hydroxybutyrylation activity (Fig. 3C). In parallel, we performed deacetylation assays with each mutant using acetylated lysine-containing fluorogenic peptides that are susceptible to fluorophore release after deacetylation. As expected, mutant mHDAC2 proteins also exhibited impaired deacetylation activity. We found a strong positive correlation between the deacetylation activity and β-hydroxybutyrylation activity (Pearson correlation coefficient r = 0.9688, Fig. 3D), indicating that the HDAC2 active site within the deacetylation domain is also responsible for Kbhb formation.

**Fig. 3.**
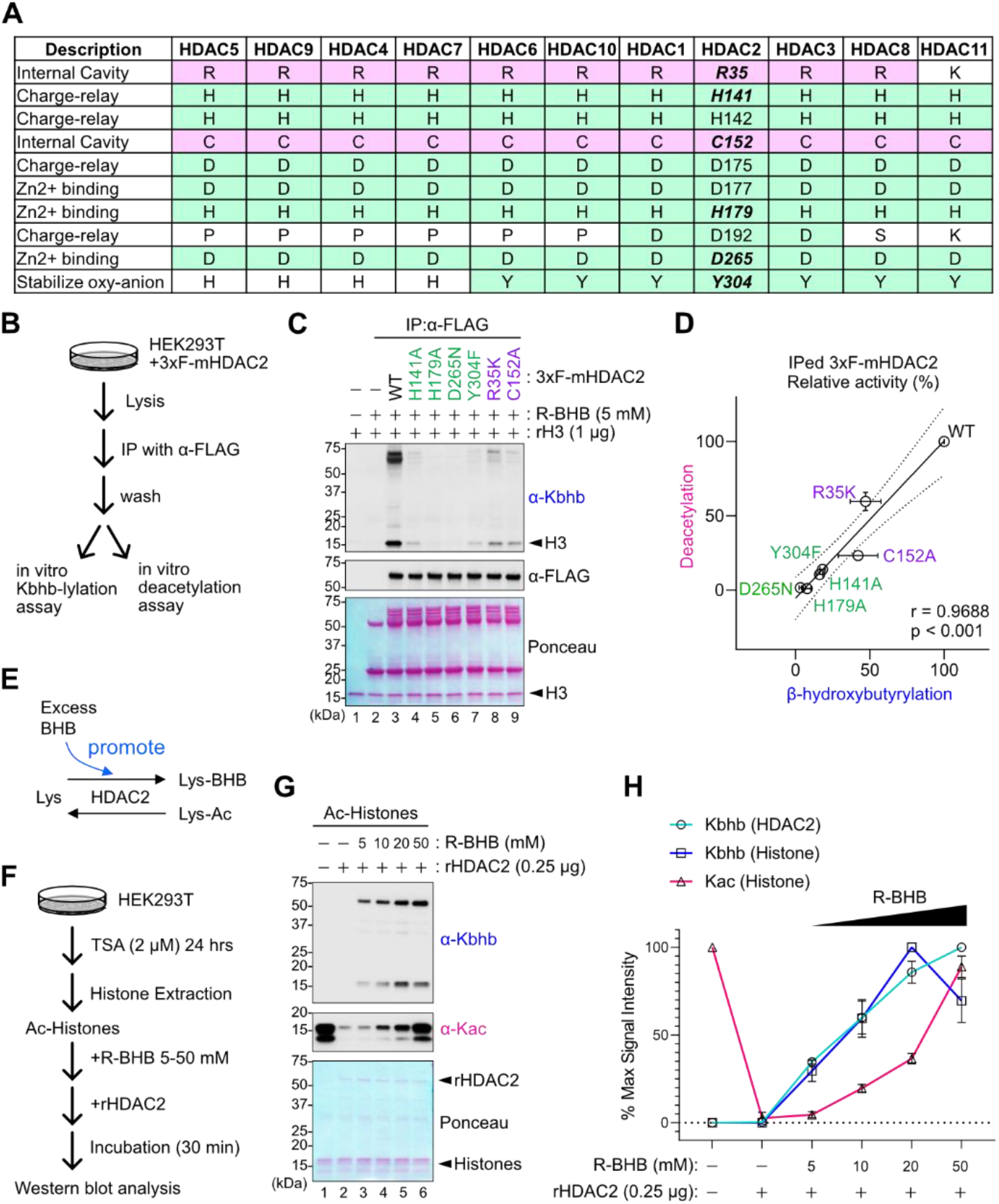
Lysine β-hydroxybutyrylation requires HDAC deacetylation residues. (**A**) Table of residues important for HDAC deacetylation activity, color-coded according to their proposed functions. The numbers in the HDAC2 column indicate the amino-acid numbers for HDAC2. The amino-acid residues with mutations are shown in italic and bold. (**B**) Schematic of experimental workflow. HDAC2 KO HEK293T cells expressing 3xFLAG-mHDAC2 were used for immunoprecipitation with α-FLAG antibody. The immunoprecipitants were used for in vitro lysine β-hydroxybutyrylation assay and in vitro deacetylation assay. (**C**) Representative western blots of in vitro lysine β-hydroxybutyrylation with IPed 3xF-mHDAC2 WT or indicated mutants. (**D**) Scatter plot of deacetylation and lysine β-hydroxybutyrylation activity for each mutant. Both activities are relative activity to WT 3xF-mHDAC2. Data are represented as mean +/-SEM of independently performed experiments. The line with 95% confidence bands (dotted line) was generated by simple linear regression. N=4 experiments for β-hydroxybutyrylation and N=3 experiments for deacetylation. Pearson correlation coefficient (r) with two-tailed p-value is indicated. (**E**) Schematic of the hypothesis that excess BHB may promote the lysine β-hydroxybutyrylation reaction. (**F-H**) Schematic of experimental workflow (F). Highly acetylated histone proteins (Ac-Histones) were isolated from TSA-treated HEK293T cells, and Ac-Histones (2.5 μg) and rHDAC2 (0.25 μg) with R-BHB at the indicated concentrations were incubated for 30 mins. The same isolated histones were pooled and used for all the three experiments. Kbhb and Kac were evaluated by western blot in N=3 independent experiments, with one representative experiment shown in (G). Relative intensities of anti-Kbhb and anti-Kac signals as compared to each maximum intensity are graphed in (H). Data are represented as mean +/-SD of N=3 independently performed experiments.

There is prior evidence that BHB interacts directly with HDACs (*4*). In 2013 Shimazu et al., showed that BHB inhibits histone deacetylation by HDACs, with IC50 = 5.3 mM for HDAC1. We speculated that such a high concentration of BHB might actually reverse HDAC activity by mass action (Fig. 3E), which could be interpreted as competitive inhibition given that Kbhb had not yet been discovered. To test this hypothesis, we isolated highly-acetylated histones by acid extraction from TSA-treated HEK293T cells and incubated them with rHDAC2 and increasing BHB concentrations in our in vitro reconstitution assay (Fig. 3F). In agreement with Shimazu et al., BHB inhibited rHDAC2 deacetylation activity in a dose-dependent manner and was most effective at exceptionally high BHB concentrations of 20-50 mM (Fig. 3G,H). Importantly, Kbhb on HDAC2 and extracted histones was detected at lower BHB concentrations and also increased with BHB concentration. These data support the previous findings and combined with our mutagenesis data, provide evidence for a model in which β-hydroxybutyrylation and deacetylation require the same active site residues and therefore may become mutually exclusive depending on substrate concentration.

## HDAC2 can catalyze lysine acylation with other short-chain fatty acids

Next, we investigated whether this new mechanism of HDAC-catalyzed protein modification would extend to other acylations. To test this, we examined whether acetate, butyrate, lactate, and propionate could be transferred to H3 in our in vitro reconstitution assay (Fig. 4A). These metabolites are present in varying concentrations in the blood (Fig. 4B) (*27-30*). Excitingly, we found that rHDAC2 can indeed catalyze lysine butyrylation (Kbu) and lactylation (Kla) under similar conditions as β-hydroxybutyrylation (Fig. 4C,D). In contrast, propionate and acetate required much higher substrate concentrations (50-100 mM) to detect respective HDAC-catalyzed lysine acylation (Fig. 4E,F). These apparent differences may be driven, in part, by different SCFA carbon lengths, and different kinetics of deacylation of each of these modifications (*17*).

**Fig. 4.**
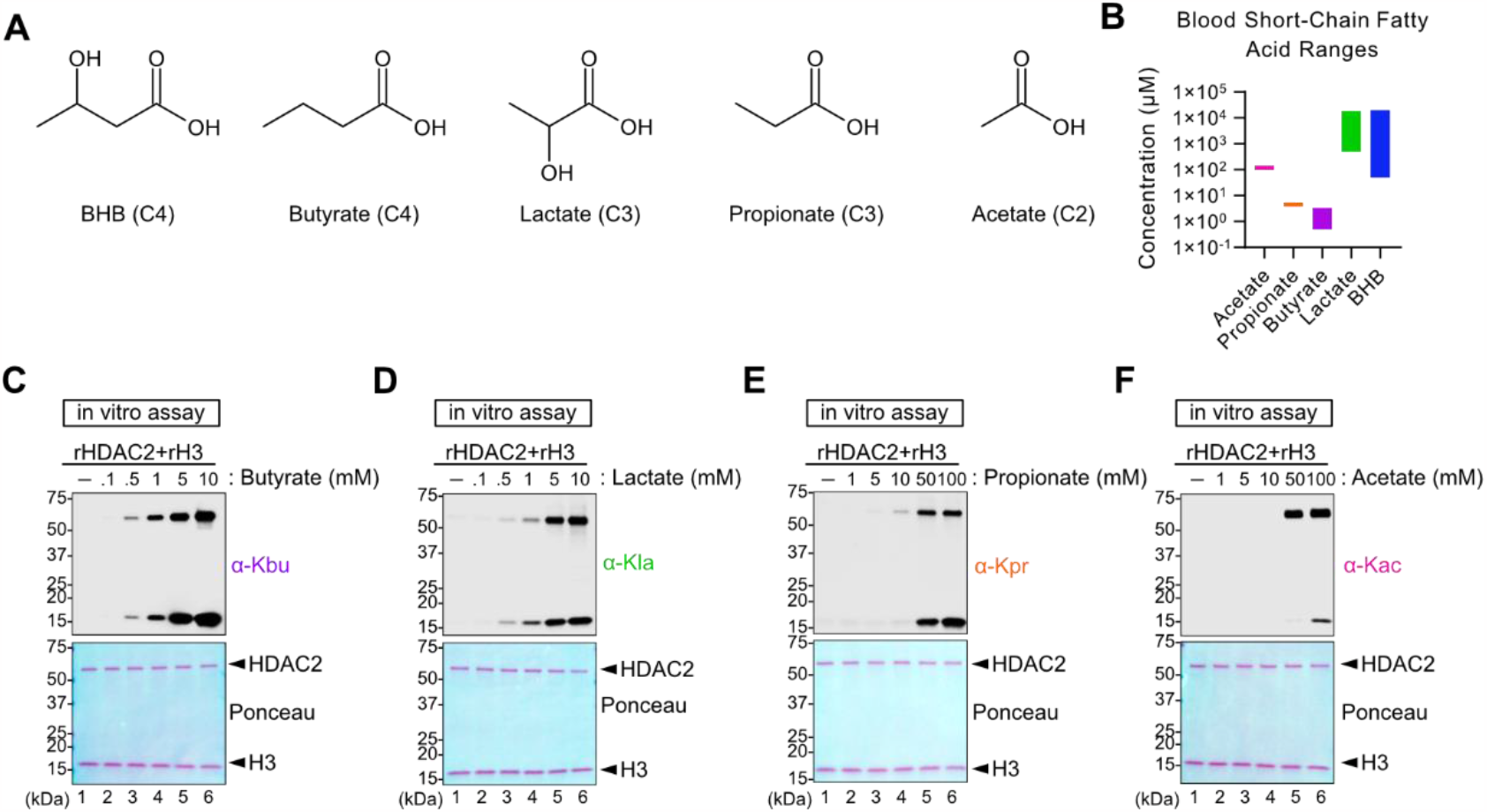
HDAC2 can acylate target proteins with other short-chain fatty acids in vitro. (**A**) Chemical structures of the indicated short-chain fatty acids are shown. (CX) next to each name indicates the number of carbons in each fatty acid. (**B**) Comparison of the concentration ranges of short-chain fatty acids in blood based on reported literature values. (**C-F**) Representative western blots of in vitro acyations by rHDAC2 with the short-chain fatty acids, Butyrate (C), Lactate (D), Propionate (E), and Acetate (F), at the indicated concentrations.

## HDAC-catalyzed Kbhb formation occurs in vivo

Finally, we tested whether HDAC-dependent β-hydroxybutyrylation could occur in animals. We addressed this question in a model of fasting-induced ketogenesis in mice. We fasted mice for 18 hours, beginning 3 hours into the overnight dark cycle when mice usually eat (Fig. 5A). The following morning, we treated male and female mice with SAHA to acutely inhibit all HDACs. Compared to vehicle-treated mice, SAHA reduced Kbhb formation in spleen (Fig. 5B-E) and bone marrow (Fig. 5F-I), with corresponding increased Kac in those organs. SAHA treatment had no major effect on Kbhb in the liver and kidney, but we also observed minimal induction of Kac in liver and kidney, suggesting the pharmacological effects of SAHA in these organs were limited (fig. S9). Altogether, our data reveal a novel function of HDACs to catalyze β-hydroxybutyrylation, and other acylations, at the molecular, cellular, and organismal levels.

**Fig. 5.**
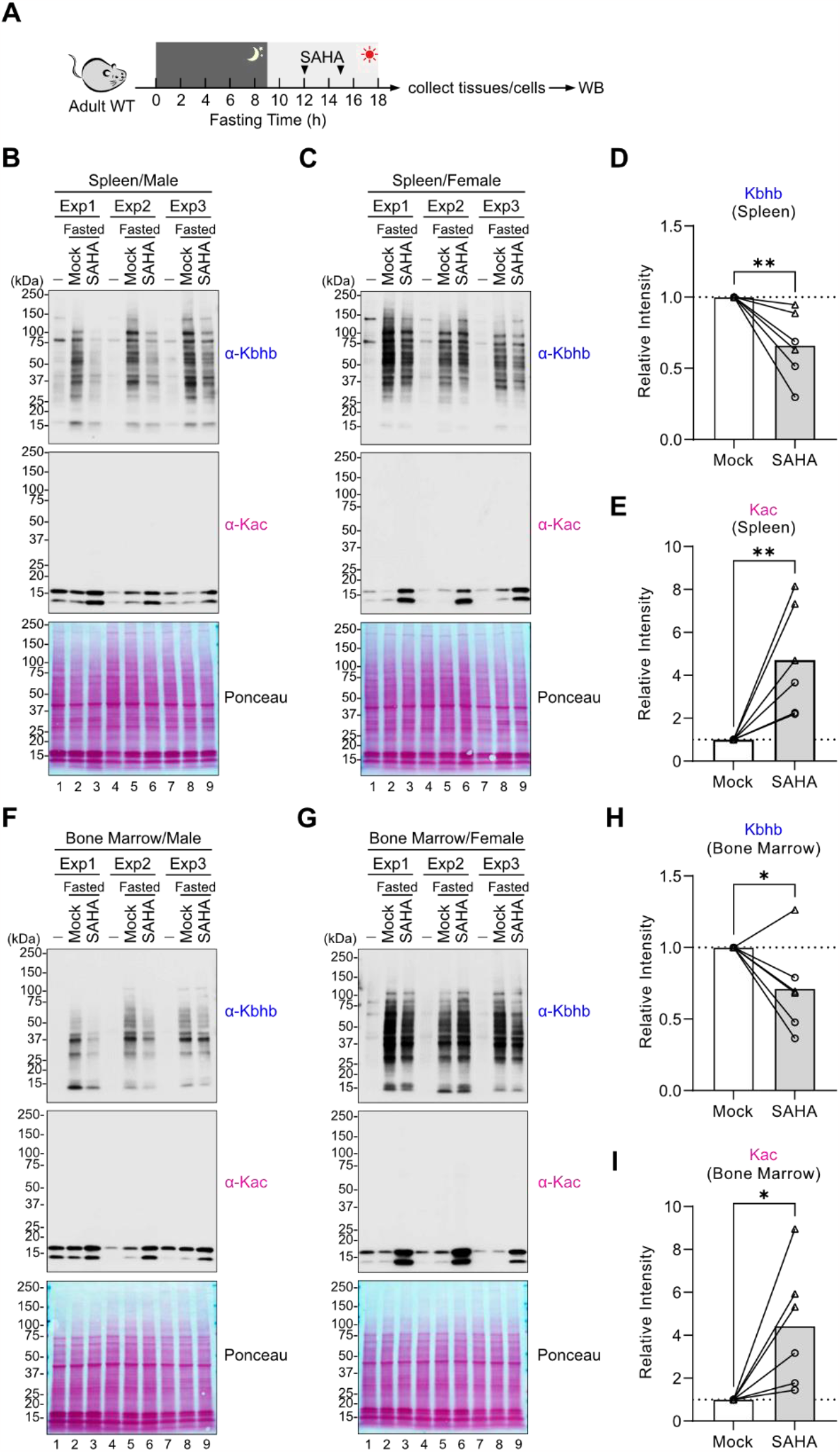
HDACs contribute to ketosis-induced Kbhb formation in mice. (**A**) Schematic of experimental design. C57BL/6 mice were fasted starting from night time and injected with SAHA twice at 12 and 15 hr after fasting was initiated. WB: western blot analysis. (**B-E**) Western blots of splenocytes are shown in (B-C). Exp1-3: N=3 independent experiments are shown on each blot, each containing a fed control, a mock-treated, and a SAHA-treated mouse as indicated. The signals were normalized by ponceau S staining and intensities relative to Mock sample for Kbhb and Kac are plotted in (D) and (E), respectively. Circles indicate males and triangles indicate females. Unpaired t-test with two-tailed P values and lines connect the control and SAHA-treated mice from the same independent experiment. (**F-I**) Western blots of bone marrow cells are shown in (F-G). Exp1-3: N=3 independent experiments are shown on each blot. The signals were normalized by ponceau S staining and intensities relative to Mock sample for Kbhb and Kac are plotted in (H) and (I), respectively. Circles indicate males and triangles indicate females. Unpaired t-test with two-tailed P values and lines connect the control and SAHA-treated mice from the same independent experiment. *p<0.05, **p<0.01

## Discussion

We report a novel regulatory mechanism of post-translational modifications in which HDACs can operate in reverse to catalyze protein acylation. Based on our mutant mHDAC2 results, combined with evidence that Kbhb abundance is proportional to BHB concentration, this is likely a consequence of shifted mass action as intracellular BHB concentrations increase. Canonical deacetylation uses hydrolysis in the HDAC active site to free acetate (or other specific acyl groups) from lysine. Here we propose that when an amenable substrate like free BHB is in the HDAC active site, the reverse condensation reaction can occur to acylate target lysine residues. Moreover, the acylating activity of class I HDACs correlates with the permissiveness of their deacylating activity across multiple short-chain fatty acids.

We suspect reverse HDAC activity was not discovered earlier because the metabolites that form more well-studied adducts do not achieve the relative changes in intracellular concentrations as compared to BHB. In support of this assumption, non-physiologically high acetate levels can also lead to HDAC2-catalyzed Kac formation in our reconstitution assay (**Fig. 4F**). In addition, canonical deacetylation rates may be fast and more enzymatically favorable, so outcompeting them could require extraordinarily high acetate concentrations that do not occur in vivo. In contrast, circulating BHB levels can increase 10-20-fold during fasting or ketogenic diet feeding when blood BHB levels can reach 2-5 mM, thus creating a wide dynamic range of substrate concentration. Our data with butyrate, lactate, and propionate suggest this principle could easily extend to other acylations that are regulated by HDACs. Future studies are needed to understand what proportion of acylations are achieved through this mechanism and how this is controlled in different metabolic conditions. Additionally, whether other protein modification enzymes are similarly reversible also has not been explored.

Understanding the impact of HDAC-catalyzed protein acylation will be an important area for future work. As deacetylation and β-hydroxybutyrylation require the same active site residues (Fig. 3), it is not yet possible to uncouple deacetylation and acylation, which would help disentangle the physiological roles of Kbhb. Currently HDAC inhibitors are used or being tested clinically in patients with cutaneous T cell lymphoma, HIV-1, and numerous other tumors/cancers. Similarly, enthusiasm surrounding potential therapeutic benefits of ketosis continues to grow so ketone supplement availability is expanding and clinical trials are ongoing in multiple diseases. Our work suggests ketone bodies may influence activity of HDAC inhibitors, or conversely, that HDAC inhibitors will also disrupt the previously unknown protein acylation activity of class I HDACs. These are potentially important diet-medication interactions to consider in disease treatment.

## Acknowledgements

We thank members of the Goldberg lab for comments and intellectual discussion. We also thank Hiten Madhani (UCSF) and Holly Ingraham (UCSF) for intellectual discussion and manuscript feedback, Jonathan Sandoval (UCSF) and Danica Galonic Fujimori (UCSF) for intellectual discussion, Anna Molofsky’s lab (UCSF) for sharing HEK293T cells, and Yoshihiro Ishikawa (UCSF) for sharing iMEFs and NIH3T3. The Goldberg lab is funded by NIH/NIA R00AG058801 to ELG, a pilot and feasibility award from the UCSF Liver Center P30DK026743 to ELG, the Chan Zuckerberg Biohub, the Sandler Program for Breakthrough Biomedical Research, which is partially funded by the Sandler Foundation (to ELG), and JSPS Overseas Research Fellowships (to TT). Dr. A. L. Burlingame’s lab is funded by the Dr. Miriam and Sheldon G. Adelson Medical Research Foundation. The Burslem lab is funded by NIGMS (R35GM142505) to G.M.B.

## Author Contributions

ELG conceptualized and oversaw project development, designed experiments, and prepared the manuscript. TT designed and performed experiments, analyzed data, and prepared manuscript. CL performed in vitro BHB-CoA experiments. MAN performed LC-MS/MS sample prep, acquisition, and analysis of in vitro reconstitution samples under the supervision of GMB. JAOP performed LC-MS/MS sample prep, acquisition, and data analysis under the supervision of ALB. BS and EB performed metabolomics sample acquisition and data analysis under the supervision of CB. All authors assisted in manuscript preparation.

## Declaration of Interests

The authors declare no competing interests.

## Notes

### Competing Interest Statement

The authors have declared no competing interest.

