## Supplemental Materials for "Reversible histone deacetylase activity catalyzes lysine acylation"

**Method**

***Cell culture***

HEK293T, iMEFs (immortalized mouse embryonic fibroblasts), NIH3T3, and U2OS cells were cultured in DMEM (Gibco, 10569044 ) supplemented with 10% fetal bovine serum (FBS) and 1x antibiotic-antimycotic (Gibco, 15240062) in a 37°C incubator with 5% $CO_2$. Cells were treated as indicated in the respective figure legends with the following reagents: DL-β-Hydroxybutyric acid sodium salt (Na-BHB, H6501, Sigma-Aldrich), (R)-3-Hydroxybutyric acid (R-BHB, 54920, Sigma-Aldrich), sodium butyrate (B5887, Sigma-Aldrich), Trichostatin A (TSA, T8552, Sigma-Aldrich), Suberoylanilide Hydroxamic Acid (SAHA, 10009929, Cayman), MS-275 (14043, Active Motif), Triacsin C (ab141888, Abcam), A485 (63875, Tocris).

***Mice***

All mice were housed and maintained in a specific pathogen-free animal facility at the University of California, San Francisco. All animal protocols were approved by the UCSF Institutional Animal Care and Use Committee. C57BL6/J were obtained from Jackson Laboratory (stock #000664). Adult male and female mice (2-4 month old) were used for experiments. 2 mg SAHA per mouse in a solution of 5% DMSO:30% PEG 300:5% Tween 80:ddH2O was administrated by intraperitoneal injection (approximately 100 mg/kg body-weight). For tissue collection, mice were euthanized by $CO_2$ and blood was collected by cardiac puncture. After perfusion with cold PBS, liver, kidney, spleen, and bone were collected. Liver and kidney were immediately snap frozen in liquid nitrogen and stored at -80°C until use. Bone marrow cells were isolated by flushing with 10% FBS/RPMI medium. Spleen was minced through a 70 µm nylon filter with 2 mL 10% RPMI medium. For both bone marrow and spleen, red blood cells were lysed with ACK lysing buffer (avantor, 118-156-101). Cells were then washed with PBS and used for protein extraction (see "Western blot analysis" section).

***siRNA transfection***

Cells were harvested after trypsinization, quenched with culture medium, and pelleted by centrifugation at 1,500 rpm for 5 mins. Cell pellet was resuspended in antibiotic-free medium. Each siRNA was directly added to each well of a 6-well dish. Subsequently, 200 µL of Opti-

MEM and 6 µL of RNAiMax (Life Technology) were directly added to the wells. After 10 mins incubation, cell suspensions ($4x10^5$ cells/well) were added to each well and antibiotic-free medium was added to bring the culture volume to 2 mL. After 24 hours, culture medium was replaced with fresh medium containing antibiotics. 24 hours later, cells were harvested and the siRNA transfection was repeated as described above. The next day, cells were treated with BHB at the indicated concentrations in each figure and collected for western blot or qPCR analysis. The following Silencer Select siRNA (Thermo Fisher Scientific) were used: s73 (HDAC1, validated), s6495 (HDAC2, validated), s16876 (HDAC3, validated), s3495 (CBP, validated), s4697 and s534247 (p300), s39121 and s39122 (ACSS1), s31745 and s31746 (ACSS2), and s35911 and s35912 (ACSS3). Each siRNA for p300 and ACSS1-3 were validated by qPCR analysis before the experiments and two siRNAs were mixed and used for efficient knockdown.

#### *Plasmids*

Mouse HDAC2 H141A cDNA was amplified from the pcDNA3.1-HDAC2-H141A (115346, Addgene), and using it as a template, mHDAC2 wild-type (WT) cDNA was generated by over-lapping PCR with primers harboring the wild-type sequence. Using the WT cDNA as a template, other mHDAC2 mutants were generated by overlapping PCR with primers harboring each mutation. These amplicons were inserted into pLenti-CMV-GFP-Puro (658-5) (Addgene #17448) using NEbuilder (NEB). Oligo DNA that contained each gRNA sequence encoding human HDAC1 or HDAC2 was inserted into lentiGuide-Puro vector (52963, Addgene) according to the Addgene resource information. The following gRNA sequences were used: g-hHDAC1#1 (CATCCGTCCAGATAACATGT), g-hHDAC1#2 (GGAGATGTTCCAGCCTAGTG), g-hHDAC2#1 (TCAAAGAGTCCATCAAACAC), g-hHDAC2#2 (CCTCCTCCAAGCATCAGTAA). Lentivirus production vectors, pMD2.G and psPAX2 were obtained from addgene. NAX_CAGGS_CAS9_Blasto is from Addgene (#167856) and the ePiggyBac transposon (PBase) vector was a kind gift from Kirstin Meyer (Orion Weiner lab, UCSF) (*1*). All plasmids used in this study were confirmed by Sanger sequencing or Nanopore sequencing (Plasmidsaurus).

#### *Lentivirus production and spinfection*

HEK293T cells were seeded at 6.6x$10^5$ cells per a well of 6-well dish. On day 2, cell culture medium was replaced with antibiotic-free medium. Cells were transfected with pMD2.G (0.80 µg), psPAX2 (0.82 µg) and each lentivirus vector (1.87 µg) using 14 µL of PEI Max (24765, Polysciences) and 170 µL of Opti-MEM, according to the manufacturer's instructions. After 24 hours, cell culture medium was replaced with fresh antibiotic-free medium and 1x ViralBoost reagent (6 µL per 3 mL culture; VB100, ALSTEM). After incubating for 48 hours and medium replacement, virus-containing culture medium was collected and centrifuged at 1,500 rpm for 5 mins to pellet any cells. The supernatant was filtered through a 0.45 µm PES filter, aliquoted into 1.5 mL tubes, and stored at -80°C until use.

For infection, cells were harvested by trypsinization, pelleted by centrifugation at 1,500 rpm for 5 mins, and resuspended at 1.5x$10^6$ cells/mL in medium containing 1 µg/mL hexadimethrine bromide (also known as polybrene; H9268, Sigma-Aldrich). The pre-determined amount of virus-containing culture (approximately 0.75x$10^6$ transduction unit) was added to each well of 12-well dish. Antibiotic-free medium containing 1 µg/mL polybrene was added to bring the final volume to 1 mL, followed by addition of the cell suspension (1 mL, 1.5x$10^6$ per well). The 12-well plate was centrifuged at 700 *xg* for 2 hours at 30°C and incubated at 37°C $CO_2$ incubator overnight. On the following day, infected cells were replated to a 6-well dish with puromycin (1 µg/mL) for drug selection.

***CRISPR/Cas9-mediated gene knockout in cells***

HEK293T cells were reverse-transfected with NAX_CAGGS_CAS9_Blasto and PBase vectors using lipofectamine 3000 (Life technology) according to the manufature's protocol. Cas9-integrated populations were selected with blastcidin at 10 µg/mL. Single cells were isolated using limited dilution to isolate Cas9-expressing HEK293T clones. Cas9+ HEK293T cells were reverse-transfected with each gRNA-expressing vector using Lipofectamine 3000 and selected with puromycin at 2 µg/mL and blastcidin at 10 µg/mL. After confirming the efficient reduction of target protein in bulk cells by western blot, single clones for each gRNA were cloned and tested.

***Western blot analysis***

Western blot analysis was performed as described previously (*2*). Cells were harvested after trypsinization and quenching with medium, washed once with PBS, and lysed with RIPA buffer (Thermo Scientific) supplemented with protease and phosphatase inhibitors (Thermo Scientific). The cell lysates were sonicated using a Branson Sonifier 450 until the viscosity of lysates disappeared and then centrifuged at 14,000 rpm (17,968 x*g*) at 4°C for 5 mins to clarify cell lysates. Protein was quantified with the DC Protein Assay Kit (BioRad Laboratories). For liver and kidney samples, frozen tissues were homogenized in RIPA buffer supplemented with protease and phosphate inhibitors by Bead Mill 24 Homogenizer (Fisher Scientific) with the following parameters: S=4.00, C=04, T=0.30 s, and D=0.10. After centrifugation at 1,500 rpm at 4°C for 5 min, the supernatants were transferred to a new 1.5 mL tube and SDS-PAGE samples for liver and kidney were then prepared as described for cells.

SDS-PAGE samples were prepared by mixing clarified lysates wth 4x Laemmli buffer/10% β-mercaptoethanol (BME) (BioRad Laboratories) and then boiled at 95°C for 5 mins. Equivalent amounts of protein were resolved by SDS-PAGE and then transferred to a nitrocellulose membrane using the Trans-Blot Turbo Transfer system (BioRad Laboratories). Membranes were stained with Ponceaus S, blocked with 5% milk/TBS-T (Tris Buffered Saline/0.1% Tween-20) for 30-60 mins, and incubated with primary antibodies, followed by secondary HRP-conjugated antibodies. Blots were developed using a chemiluminescent substrate (SuperSignal™ West Pico PLUS or SuperSignal™ West Femto Maximum Sensitivity Substrate; Thermo Scientific) and imaged with an Azure 300 (Azure Biosystems). Ponceau S staining was used to confirm equal total protein loading across samples.

***Antibodies***

The following antibodies were used in this study: anti-Kbhb (monoclonal, PTM Biolabs, PTM-1201RM) which is mainly used in this study, anti-Kbhb (polyclonal, PTM Biolabs, PTM-1201) which is only used in Figure S1, anti-Kac antibody mix (CST, 9814S), anti-Kbu (PTM Biolabs, PTM-301), anti-Kpr (PTM Biolabs, PTM-203), anti-Kla (PTM Biolabs, PTM-1401RM), anti-HDAC1 (CST, 5356), anti-HDAC2 (CST, 5113), anti-HDAC3 (CST, 3949), anti-FLAG (Sigma-aldrich, F3165), Goat anti-Rabbit IgG HRP (Thermo Scientific, 31460), Goat anti-Mouse IgG HRP (Thermo Scientific, 62-6520), Mouse TrueBlot® ULTRA: Anti-Mouse Ig HRP (ROCKLAND, 18-8817-31), and Rabbit TrueBlot®: Anti-Rabbit IgG HRP (ROCKLAND, 18-8816-31).

#### *RNA extraction and qPCR analysis*

Total RNA was isolated using RNeasy micro kit (Qiagen, 74034) according to the manufacturer's protocol. cDNA was generated from equal amounts of RNA using iScript cDNA synthesis kit (Bio-rad, 1708891). Quantitative PCR (qPCR) with Power SYBR™ Green PCR Master Mix (Applied Biosystems™, 4367659) was carried out using CFX384 real-time PCR detection system (Bio-rad). The primer sequences are provided in Table S3.

#### *Sample preparation for targeted metabolomic analysis*

Cells were seeded at a density of $1.0 \times 10^6$ cells/well in a 6-well dish. Six wells were used as technical replicates for each condition. After one day of culture, BHB, butyrate, or BHB+butyrate, all at 5 mM, was directly added to the culture medium. After 24 hours, culture medium was removed and 0.9% NaCl was added. Cells were harvested by scraping, and $1.0 \times 10^6$ cells were transferred to a collection tube (2 mL polypropylene screw top vials). After centrifugation at 1,500 rpm to remove the supernatant, each cell pellet was resuspended in 0.4 mL ice-cold LC-MS grade methanol, and tubes were immediately placed on ice. After a 5 minute incubation, 0.4 mL ice-cold LC-MS grade water was added to each tube, and samples were stored at -80°C until further processing.

#### *Development of Butyryl-CoA and BHB-CoA multiple reaction monitoring signals*

Multiple reaction monitoring ion pairs were established for butyryl-CoA and BHB-CoA from synthetic standards at method appropriate flow rate and buffer conditions on a Sciex 5500 QTRAP® mass spectrometer. For butyryl-CoA the following daughter ions were used for negative mode detection in combination with the parent mass (836.0): 489.0, 426.0, 408.0, 79.0. For BHB-CoA the following daughter ions were used in negative mode in combination with the parent mass (852.0): 505.0, 426.0, 408.0, 79.0. Resolution of BHB-CoA from isobaric malonyl-CoA was ensured chromatographically (*3*).

#### *Metabolite and Lipid Sample Preparation*

For all LCMS methods, LCMS grade solvents (Thermo Fisher Scientific) were used. Samples were collected in 0.4 mL of ice-cold methanol followed by 0.4 mL of water. Cells were scraped to recover insoluble products and transferred to a microtube. To each sample, 0.4 mL of chloroform was added. Samples were agitated for 30 minutes at 4°C and centrifuged at 16,000 xg for 20 min. 400 µL of the top (aqueous) layer was collected. A subaliquot of the aqueous layer was taken for O-benzylhydroxylamine (O-BHA) derivatization of carboxylic acids. The aqueous layer was diluted 5x in 50% methanol in water and prepared for LCMS injection.

#### *Short Chain Fatty Acid Derivatization*

To reliably capture butyrate and increase sensitivity for BHB, samples were derivatized O-BHA according to previously established protocols with modifications (*4, 5*). An aqueous pyridine buffer at 1M pyridine and 0.5 M hydrochloric acid was prepared fresh. A volume of 35 µL of the aqueous metabolite extract was sub-aliquoted and 10 µL of 1M O-BHA in reaction buffer and 10 µL of 1M 1-Ethyl-3-(3-dimethylaminopropyl)carbodiimide in reaction buffer were added. Samples were shaken at room temperature for 2 hrs. The reaction was quenched with 50 µL of 0.1% formic acid. Derivatized carboxylic acid compounds were extracted via the addition of 400 µL ethyl acetate. Following mixing and centrifugation 16,000 *xg* for 5 min at 4°C to induce layering, the upper (organic) layer was collected and taken to dryness under vacuum. Samples were resuspended in 300 µL of water for LCMS injection.

#### *Liquid Chromatography Mass Spectrometry*

Tributylamine was purchased from Millipore Sigma. All samples were separated using a Sciex ExionLC™ AC system and measured using a Sciex 5500 QTRAP® mass spectrometer.

For detection of acyl-CoA conjugates signals were collected as part of a method looking at a series of central metabolic metabolites based on previously established methods (*6*). Samples were injected onto a Waters™ Atlantis T3 column (100Å, 3 µm, 3 mm X 100 mm) and eluted using a binary gradient from 5 mM tributylamine, 5 mM acetic acid in 2% isopropanol, 5% methanol, 93% water (v/v) to 100% isopropanol over 15 minutes. Peaks were compared to synthetic standards retention time and ratios between MRMs.

O-BHA-derivatized samples were separated with a Waters™ Atlantis dC18 column (100Å, 3 µm, 3 mm X 100 mm) and eluted using a 6 min gradient from 5-80 % B with buffer A as 0.1 % formic acid in water and B as 0.1 % formic acid in methanol. Butyrate and BHB were detected using MRMs from previously established methods and identity was confirmed by comparison to derivatized standards (*4, 5*).

All signals were integrated using MultiQuant® Software 3.0.3. Acyl-CoA's were quantified via the 79.0 phosphate MRM after signal confirmation with all four MRMs for each molecule. Signals were normalized to the total signal across all collected metabolite signals to correct for different sample starting abundances prior to statistical comparison between treatments.

#### *Immunoprecipitation (IP) for Kbhb proteome*

HEK293T cells on 10-cm dish (3 dishes) were treated with 10 mM Na-D/L-BHB for 3 hours, and then harvested by trypsinization and quenching with medium, and washed once with PBS. Approximately $1.0 \times 10^7$ cells per tube were lysed with 150 µL SDS-lysis buffer (50 mM Tris-HCl at pH7.5, 1% SDS). The cell lysates were sonicated using a Branson Sonifier 450 and protein concentration was measured with DC Protein Assay Kit (BioRad Laboratories). After adding BME at a final concentration of 0.5% to approximately 16 mg/mL lysates, samples were boiled at 95°C for 5 mins to denature proteins. The denatured proteins were diluted 1:10 with NP-40 lysis buffer (50 mM Tris-HCl at pH 7.5, 150 mM NaCl, 1 mM EDTA, 10% glycerol, 1% Nonidet P-40) to reduce SDS concentration. The approximately 1.6 mg/mL lysates (700 µL per sample) were incubated with antibody-conjugated ProteinA/G Dynabeads (Invitrogen, 88802) overnight at 4°C. Three independent control (Normal rabbit IgG, 2729S, CST) and three anti-Kbhb (PTM biolabs, PTM-1201RM) samples were processed in parallel. Resin was washed five times with NP-40 lysis buffer, followed by five washes with 50 mM ABC buffer (0.4% Ammonium Bicarbonate/dH2O), and eluted by incubation in 60 µL of 0.2% RapiGest-elution buffer (50 mM ABC, 0.2% RapiGest SF from Waters) at 50°C for 20 mins. A portion of each elution ($1/6^{th}$) was mixed with 4x Laemmli buffer/10% BME (BioRad Laboratories) and boiled at 95°C for 5 minutes for western blot analysis. The remainder of each elution was stored at -80°C until mass spectrometry analysis.

#### *Mass spectrometry analysis (Kbhb proteome)*

Protein was reduced by adding 5 mM DTT and incubating at 60°C for 30 min. After that, samples were treated with 10 mM iodoacetamide at room temperature for 30 min and digested with 500 ng trypsin (mass spectrometry grade, Promega) for 16 h at 37°C temperature. After this, another 500 ng aliquot of the digestion enzyme was added, and the digestion was allowed to continue for 4 h at 37C. Samples were added 5% formic acid and incubated 37°C for 30 min.

Digested material was recovered using C18 ZipTips (Waters), eluted in 50% acetonitrile 0.1% formic acid, evaporated and resuspended in 0.1% formic acid for mass spectrometry analysis. Peptide mixtures were loaded onto a 2 um 75um x 50 cm PepMap RSLC C18 EasySpray column (Thermo Scientific). 3-hour water/acetonitrile gradients (2–25% in 0.1% formic acid) were used to elute peptides, at a flow rate of 200 nl/min, for analysis in a Orbitrap Lumos Fusion (Thermo Scientific). MS analysis was performed in positive ion mode. MS spectra were acquired between 375 and 1500 m/z with a resolution of 120000. For each MS spectrum, multiply charged ions ions over the selected threshold (2E4) were selected for MSMS in cycles of 3 seconds with an isolation window of 1.6 m/z. Precursor ions were fragmented by HCD using a relative collision energy of 30. MSMS spectra were acquired in centroid mode with resolution 30000 from m/z=110. A dynamic exclusion window was applied which prevented the same m/z from being selected for 30s after its acquisition.

For peptide and protein identification, peak lists were generated using PAVA in-house software (*7*). All generated peak lists were searched against the human subset of the SwissProt database (SwissProt.2019.07.31), using Protein Prospector (*8*) with the following parameters: Enzyme specificity was set to Trypsin, and up to 2 missed cleavages per peptide were allowed. Carbamidomethylation of cysteine residues, was allowed as fixed modification. N-acetylation of the N-terminus of the protein, loss of protein N-terminal methionine, pyroglutamate formation from of peptide N-terminal glutamines, oxidation of methionine were allowed as variable modifications. Mass tolerance was 10 ppm in MS and 30 ppm in MS/MS. The false positive rate was estimated by searching the data using a concatenated database which contains the original SwissProt database, as well as a version of each original entry where the sequence has been randomized. A 1% FDR was permitted at the protein and peptide level.

#### *Downstream analyses for Kbhb proteome*

Obtained protein (peptide) list was further analyzed by SAINT (Significance Analysis of INTeractome) analysis in MAC-tag PPI (http://proteomics.fi/) (*9*). We considered proteins with

SAINT score > 0.66 and log2(Fold Change; FC, Kbhb vs Mock) >1 as Kbhb-IP-enriched proteins. This SAINT score corresponds to an estimated protein-level Bayesian false discovery rate (FDR) of <0.05. The SAINT score and log2FC plot was generated using “ggplot2” package in R (ver. 4.2.2). Using the “clusterProfiler” package in R, we performed redundancy-reduced gene ontology (GO) analysis on the Kbhb-enriched-protein list and used the top 20 enriched GO terms for biological processes and cellular compartment for visualization. Localization information for our Kbhb proteome was obtained from the human protein atlas (https://www.proteinatlas.org/) whenever possible and counted in R. We only considered “Subcellular.main.location” but ignored “Subcellular.additional.location” for this analysis. The nested pie chart for the localizations was generated in Excel (Microsoft) and edited in Affinity Designer 2 (Affinity). Two subcellular localizations “Midbody ring” and “Mitotic spindle” were too few (only 1 protein for each) to be shown on the chart so they are not depicted despite being included in the analysis. We used STRING analysis on our Kbhb proteome, and HDAC1/2-realted proteins (“physical direct association”) were identified based on the “experimental evidences and database”. STRING analysis (https://string-db.org/) for the HDAC1/2-related proteins was re-run, and the network was visualized. The resulting image was slightly edited in Affinity Designer 2 (Affinity) to change font sizes and colors.

***Sequence alignment and phylogenic tree generation for HDACs***

All human HDAC sequences were retrieved from Uniprot (https://www.uniprot.org/) and aligned by CLUSTAL W with the default setting in MEGA11 (*10*). A phylogeny tree was created with the alignment using the Maximum likelihood method with the Bootstrap test (n=500) in MEGA11.

***In vitro acylation assay with recombinant HDACs***

Most in vitro reconstitution experiments used recombinant proteins from Cayman: rHDAC1 (10009231), rHDAC2 (36419), rHDAC3/NCOR2 (10009232), rHDAC4 (10009652), rHDAC5 (10009379), rHDAC6 (10009465), rHDAC8 (19380). rHDAC2 protein from Activ Motif (31505) was used for experiments shown in Fig. S6, and rHDAC2 from BPS Bioscience (50002) was used for Fig. 2G-I and Fig. S7. We selected vendors based on the availability of recombinant proteins from each company at that time and observed no difference in rHDAC2 acylation activity among the vendors. Recombinant Histone H3.1 (rH3) was from NEB (M2503S). In most experiments, 1 µg rHDAC and/or 1 µg rHDAC  was incubated in reaction buffer (50 mM Tris-HCl pH 7.5) with either (R)-3-Hydroxybutyric acid (R-BHB) (54920, Sigma-Aldrich), DL-β-Hydroxybutyric acid sodium salt (Na-BHB) (H6501, Sigma-Aldrich), sodium butyrate  (B5887, Sigma-Aldrich), sodium acetate (S2889, Sigma-Aldrich), sodium propionate (P1880, Sigma-Aldrich) or sodium L-lactate (71718, Sigma-Aldrich) for the indicated time (typically 30-60 mins) at 37°C. In the pH test experiments, 50 mM HEPES-KOH was used as the reaction buffer instead of 50 mM Tris-HCl; no difference in β-hydroxybutyrlyation activity was observed in HEPES versus Tris buffers. Reactions were performed in 8-strip tubes in a thermal cycler (ProFlex PCR system, applied biosystems) in a final reaction volume of 15 µL. Reactions were stopped by adding 4x Laemmli buffer or acetic acid (final. 0.15%) for western blot or mass-spectrometric analysis, respectively.

***Mass spectrometry analysis for in vitro acylated proteins***

For propionylation equal amounts of protein were taken across the samples. Samples were dried under vacuum via SpeedVac before being resuspended in 30 µl of 50 mM of tri-ethyl ammonium bicarbonate buffer (TEABC) pH 8.5. To this solution was added 12 µl of propionylation reaction mixture (propionic anhydride and isopropanol in a 1:3 ratio) and the pH adjusted to pH 8 with NH4OH. The reaction was incubated at 37°C for 15 minutes, concentrated to dryness by SpeedVac and repeated twice. Finally, the samples were dried under vacuum via SpeedVac.

Histones were digested using Trypsin/Lys-C protease mix in 30:1 enzyme: protein ratio. Vacuum dried samples were resuspended in 30 µl of 50mM (TEABC) pH 8.5, pH was adjusted to 8.5 by 1.5 M Tris buffer. Digestion was performed by incubating samples at 37°C overnight. The reaction was stopped by adding 0.1% formic acid. Digested peptides were concentrated to dryness by SpeedVac, reconstituted in 0.1% formic acid and desalted using C18 Stage-Tips.

The C18 cleaned peptides were analyzed on Thermo Scientific Orbitrap Exploris 240 mass spectrometer interfaced with Thermo Scientific UltiMate 3000 HPLC and UHPLC Systems. Peptide digests were reconstituted in 0.1% formic acid and were separated on an analytical column (75 µm × 15 cm) at a flow rate of 300 nL/min using a step gradient of 1–25% solvent B (0.1% formic acid in 100 % acetonitrile) for the first 100 minutes and 25–30% for next 12 minutes, 30-70% for 4 minutes, 70–1% for next 4 minutes the total run time was set to 120 min. The mass spectrometer was operated in data-dependent acquisition mode. A survey full scan MS (from m/z 400–1600) was acquired in the Orbitrap with a resolution of 6000. Data were acquired in topN with 13 dependent scans. Fragmented using normalized collision energy with 35 % and detected at a mass resolution of 60,000. Dynamic exclusion was set for 8 s with a 10 ppm mass window.

MS/MS searches were carried out using SEQUEST search algorithms against a custom made database for human histones using Proteome Discoverer (Version 3.0, Thermo Fisher Scientific, Bremen, Germany). The workflow included Spectrum files, Spectrum selector, SEQUEST search nodes, Target decoy PSM validator, IMP-ptmRS, peptide validator, event detector, precursor quantifier. Oxidation of methionine, propionylation at lysine, beta-hydroxybutyrylation at lysine and N-terminal protein acetylation were used as dynamic modifications. MS and MS/MS mass tolerances were set to 10 ppm and 0.05 Da, respectively. Trypsin/Lys-C was specified as proteases and a maximum of two missed cleavage was allowed. Target-decoy database searches used for calculation of false discovery rate (FDR) and for peptide identification FDR was set at 1 %. Feature mapper and precursor ion quantifier were used for label-free quantification.

***FLAG immunoprecipitation followed by In vitro acylation and in vitro deacetylation assay***

HEK293T cells (80-90% confluent) on a 10-cm dish (1 dish per sample) were harvested by trypsinization and quenching with medium and washed once with PBS. Cells were lysed with 1000 µL High-Salt NP-40 lysis buffer (50 mM Tris-HCl at pH 7.5, 500 mM NaCl, 1 mM EDTA, 10% glycerol, 1% Nonidet P-40, 1x Halt Protease Inhibitors). The cell lysates were sonicated using a Branson Sonifier 450 and clarified by centrifugation at 14,000 rpm for 5 minutes. The clarified lysates were incubated with 15 µL FLAG M2 beads (M8823, Sigma) for 2 hrs at 4°C. The resins were then washed four times with the High-Salt NP-40 lysis buffer, followed by two additional washes with 50 mM Tris-HCl pH 7.5, and resuspended in 300 µL Tris-HCl pH7.5. The 200 µL of IPed samples was used for in vitro lysine β-hydroxybutyrylation reconstitution assay and the remainder (50 µL per well, duplicate) was used for in vitro deacetylation assays.

For β-hydroxybutyrylation assays, IP'ed beads containing immobilized HDAC2 were transferred to a 8-strip tube and the buffer was completely removed. All the reagents other than rHDAC2 was added to the 8-strip tubes and in vitro acylation was performed as described above. For in vitro deacetylation assay, IP'ed beads were transferred to a 96-well half area assay plate (3994, CORNING) and the buffer was completely removed. Deacetylation assay with fluorescent molecule-conjugated peptide was performed using HDAC Fluorometric Activity Assay Kit (10011563, Cayman), according to the manufacture's protocol. The reaction volume was 85 µL and deacetylation reaction was performed for 30 mins. The fluorescent signals in duplicates were detected by plate reader at excitation=350nm and emission=460nm (SpectraMax iD5, Molecular Devices). Relative IP'ed 3xFLAG-mHDAC2 amounts were estimated based on in-parallel anti-FLAG western blot analysis and used for the normalization toward both in vitro lysine β-hydroxybutyrylation activity and in vitro deacetylation activity.

#### *FLAG immunoprecipitation followed by western blot analysis*

3xFLAG-mHDAC2 was immunoprecipitated as desceibed above with the modification of using 10 µL FLAG M2 beads and Normal-Salt NP-40 lysis buffer (50 mM Tris-HCl at pH 7.5, 150 mM NaCl, 1 mM EDTA, 10% glycerol, 1% Nonidet P-40, 1x Halt Protease Inhibitors). IPed proteins were eluted in 10 µL SDS-Elution buffer (50 mM Tris-HCl at pH 7.5, 0.2% SDS) by incubating at 60°C for 10 mins. Eluted proteins were further processed by adding 5 µL 4x Laemmli SDS sample buffer and immediately boiling at 95°C for 5 mins. Proteins were analyzed by western blotting.

#### *Preparation of inactivated whole cell lysates and high acetylated histones from HEK293T cells for in vitro acylation*

For whole cell lysates, HEK293T cells were harvested and washed with PBS. Approximately $1.0x10^6$ cells were lysed with 100 uL of Buffer 1 (50 mM Tris-HCl pH 7.5, 300 mM NaCl, 5 mM MgCl2, 0.5 mM ZnCl2, 1 mM DTT, 1xHalt Protease inhibitor). Cell lysates were sonicated using a Branson Sonifier 450, incubated at 60°C for 30 mins to inactivate endogenous HDACs, and centrifuged at 14,000 rpm for 5 min to pellet the insoluble proteins. Protein content in the supernatants (soluble protein fraction) were quantified with the DC Protein Assay Kit (BioRad Laboratories), and 5 µL of the inactivated whole lysates were used for in vitro acylation assays.

To generate highly acetylated histones, we treated HEK293T cells on a 10-cm dish (approximately 50% confluency) with 2 µM TSA overnight. Cells were harvested and washed with PBS once, and $5.0x10^6$ cells were transferred to a new 1.5 mL tube. Cell pellets were lysed in 1 mL of PBS-T (1% Triton-X/PBS/1xHalt protease inhibitor) on ice for 10 mins. After centrifugation at 10,000 rpm for 1 min at 4°C, supernatant was completely removed and the pellet was washed in PBS-T again. Cell pellets were resuspended in 200 µL of extraction buffer (0.5N HCl/10% glycerol in dH2O) and incubated on ice for 30 mins. After centrifugation at 14,000 rpm for 5 min at 4°C, the supernatant was transferred to a new 1.5 mL tube and mixed with 200 uL of 1M Tris Base (pH not adjusted) to neutrize pH. The extraction buffer was replaced with histone storage buffer (50 mM Tris-HCl pH 7.5, 300 mM NaCl, 1 mM EDTA, 1 mM DTT) using an Amicon filter tube (10 kDa, 0.5 mL; UFC501024, Millipore). Final protein concentration was measured with the DC Protein Assay Kit (BioRad Laboratories) and approximately 2.5 µg extracted histones and 0.25 µg rHDAC2 were used for the assays.

***In vitro acylation with acyl-CoA***

For cell lysates, HEK293T cells were harvested and washed with PBS. Approximately $5.0x10^6$ cells were lysed with 500 uL of Buffer 2 (50 mM Tris-HCl pH 8.0, 150 mM NaCl, 1xHalt Protease inhibitor). Buffer 2 with a pH of 7.5 was also used for BHB-CoA. Cell lysates were sonicated using a Branson Sonifier 450, and centrifuged at 14,000 rpm for 5 min to pellet the insoluble proteins. The lysates were incubated with acetyl-CoA (A2141, Sigma-Aldrich), BHB-CoA (H0261, Sigma-Aldrich), or butyryl-CoA (B1508, Sigma-Aldrich) at 1 mM or 5 mM at 37°C for 2 hours.

For purified histone H3.1 (M2503S, NEB) or bovine serum albumin (BSA, A9647, Sigma-Aldrich), each protein was incubated in Buffer 2 (with the indicated pH) with either BHB (200 µM), BHB + CoA (200 µM, C3144, Sigma-Aldrich), or BHB-CoA (500 µM) at 37°C for 2 hours.

***Stastical analyses***

Statistical analyses were performed in GraphPad Prism (version 10) unless otherwise indicated. For comparison of two groups, unpaired t-test was used. For comparison of multiple test conditions to a control group, one-way analysis of variance (ANOVA) with post-hoc

Dunnett's test was used. For multiple comparisons between all groups, one-way ANOVA with post-hoc Tukey's test was used. For multiple comparisons involving two factors, two-way ANOVA with post-hoc Sidak's test was used. Correlation analysis was performed using linear regression. Analysis information for each figure is indicated in the respective figure legends. For all experiments, $p < 0.05$ was considered significant. *p<0.05, **p<0.01, ***p<0.001, ****p<0.0001. We performed experiments at least twice independently to confirm reproducibility of the results shown in this study.

### Supplemental Figures

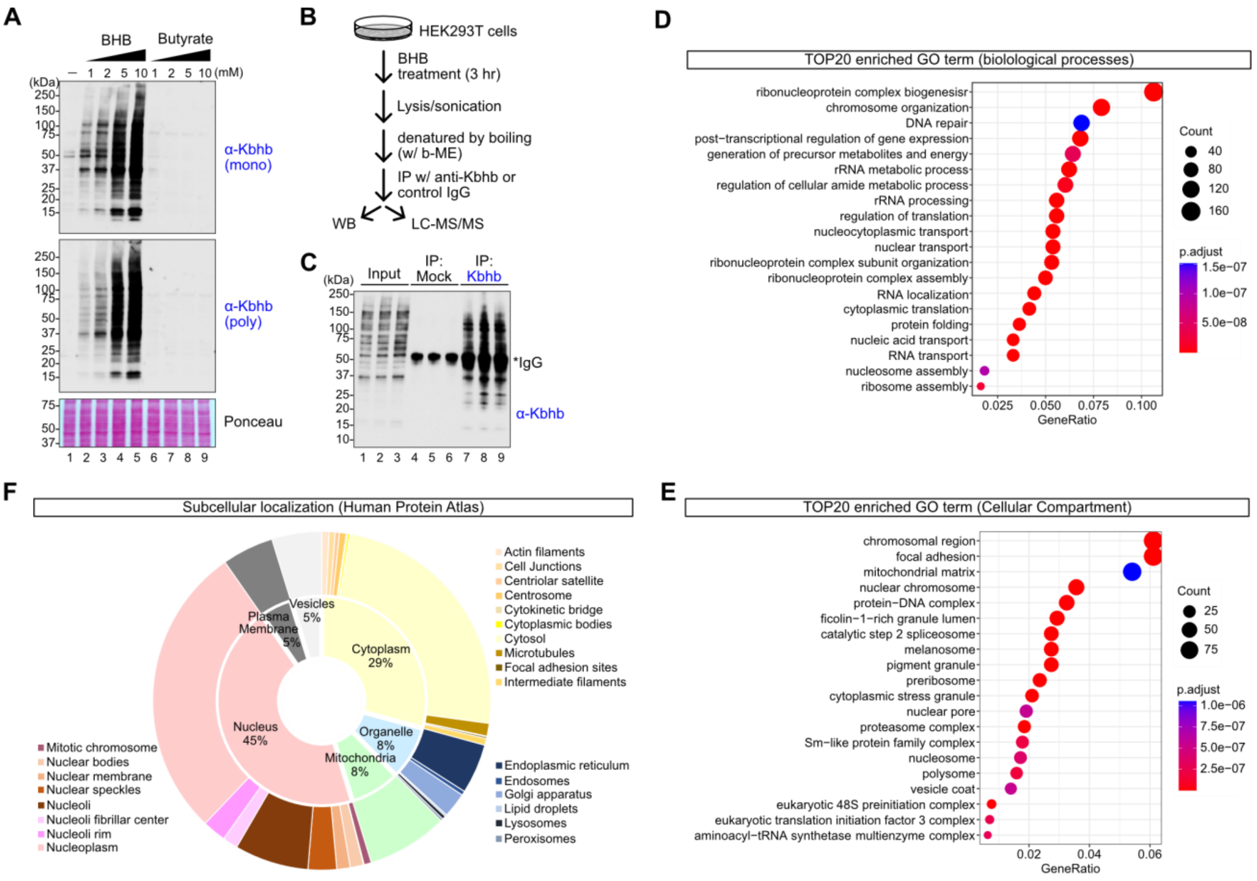


**Figure S1. Features of the Kbhb proteome**

**(A)** Representative western blots of HEK293T cells treated with various concentrations of BHB or butyrate for 24 hours. We obtained similar results with both monoclonal and polyclonal Kbhb antibodies and therefore used the monoclonal antibody for subsequent analyses. **(B)** Schematic of experimental workflow. HEK293T cells were treated with 10 mM BHB for 3 hours. The cell lysates were denatured by heating and used for immunoprecipitation with α-Kbhb antibody or control rabbit IgG. The elutions from immunoprecipitants were used for western blot and LC-MS/MS analysis. **(C)** Western blot of input and IPed samples. Each lane corresponds to an individual technical replicate that was used for mass spectrometry analysis. **(D-E)** TOP20 enriched gene ontology (GO) terms for biological processes (D) and cellular compartment (E). **(F)** Distribution of subcellular localizations for the Kbhb proteome are shown as a nested pie chart. Each protein localization was retrieved from the human protein atlas. The inner pie chart indicates larger classification, and the outer pie chart shows a finer classification.

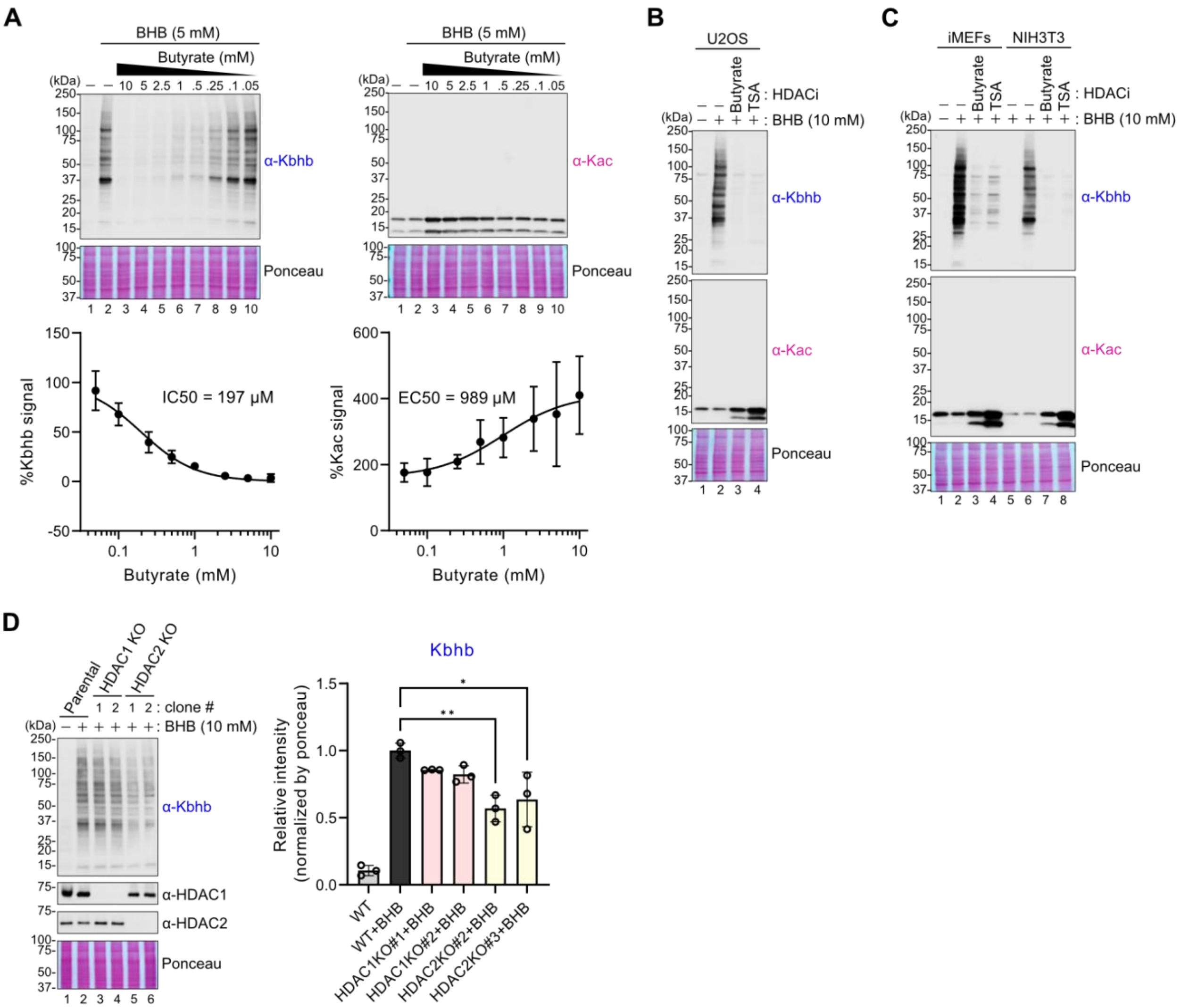


**Figure S2. HDAC inhibition blocks Kbhb formation**

**(A)** Top: Representative western blots of Kbhb and Kac abundance in HEK293T cells treated with 5 mM BHB together with butyrate at the indicated concentrations for 24 hours. Bottom: Relative intensities of anti-Kbhb (left) or anti-Kac (right) signals as compared to lane 2 (5 mM BHB treatment). The signals were normalized by ponceau S staining. Data are represented as mean +/- SEM of N=3 independently performed experiments. IC50 or EC50 were determined by nonlinear regression (curve fit). **(B-C)** Representative western blots of Kbhb and Kac abundance in U2OS (human), iMEFs (mouse), and NIH3T3 (mouse) cells treated with 10 mM BHB together with butyrate (5 mM) or TSA (2 μM) for 24 hours. **(D)** Left: Representative western blots of Kbhb, HDAC1, and HDAC2 expression in HDAC1-KO or HDAC2-KO HEK293T cells treated with 10 mM BHB for 3 hours. Right: Relative intensities of anti-Kbhb signals as compared to lane 2 (10mM BHB treatment of parental cells). The signals were normalized by ponceau S staining. Data are represented as mean +/- SD of technical triplicates. Statistical differences were calculated by 1-way ANOVA followed by Dunnett’s test for multiple comparisons. *$p<0.05$, **$p<0.01$.

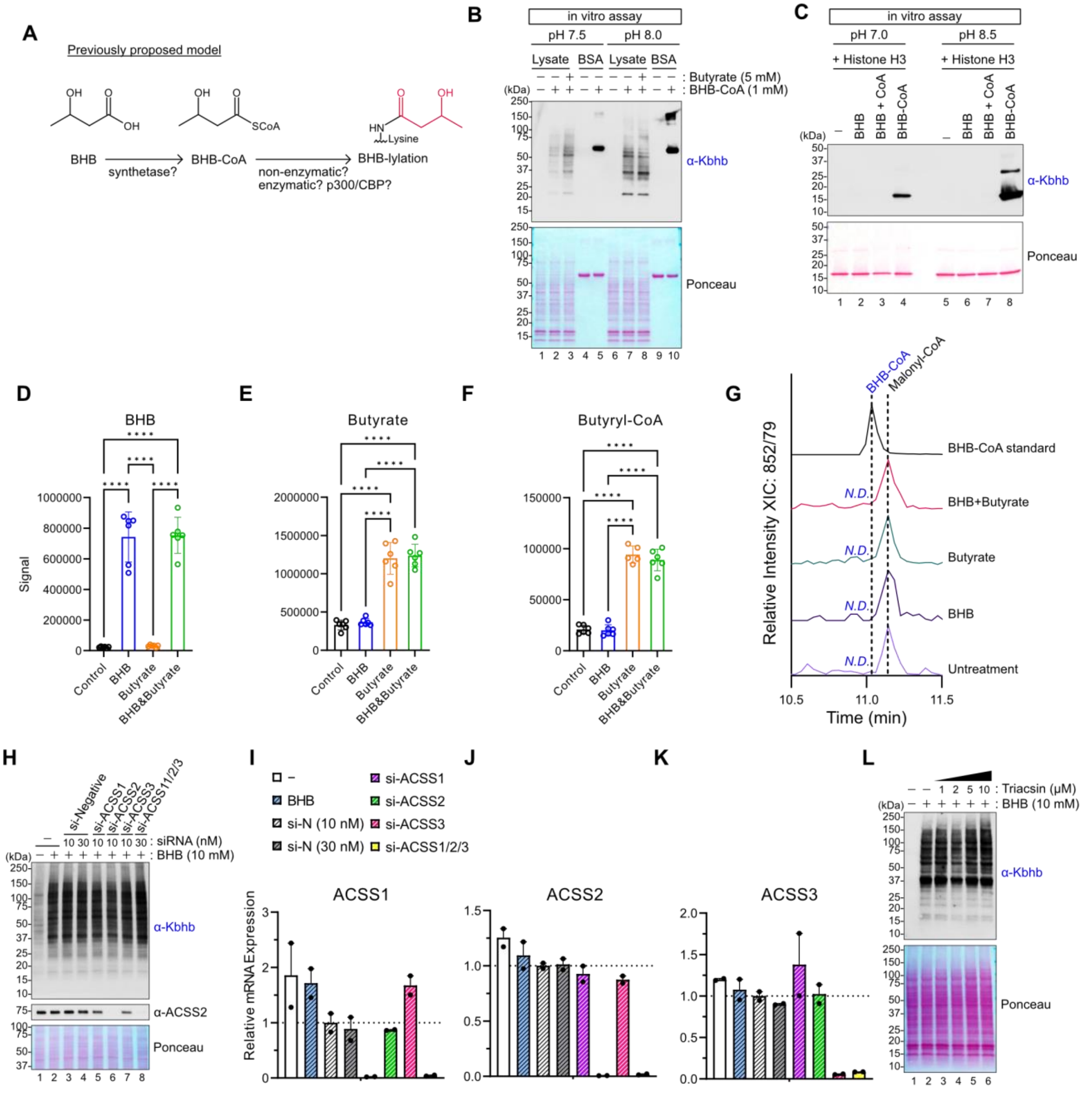


**Figure S3. Kbhb formation does not require enzymatic production or transfer of BHB-CoA**

**(A)** Schematic of the previousy proposed model for Kbhb formation. BHB was thought to be converted to BHB-CoA, which can directly modify lysines via non-enzymatic reaction or can be utilized for enzyme-dependent lysine β-hydroxybutyrylation. CBP/p300 have been proposed to be lysine BHB-transferases. **(B)** Representative western blots evaluating Kbhb formation from in vitro assay with pre-formed BHB-CoA. BHB-CoA was incubated with HEK293T cell lysates or bovine serum albumin (BSA) in the Tris-HCl buffer at the indicated pH. Butyrate was added to the reaction buffer to see if butyrate affects BHB-CoA-dependent Kbhb formation. **(C)** Kbhb formation was assessed by western blot. Pre-formed BHB-CoA or individual components BHB+CoA were incubated with recombinant histone H3 in the Tris-HCl buffer at the indicated pH. **(D-G)** HEK293T cells were treated with 5 mM BHB, 5 mM butyrate or both. After 24 hours of treatment, cellular metabolites were extracted and subjected to targeted metabolomics by LC-MS/MS. Relative intensities of BHB (D), butyrate (E), and butyryl-CoA (F) in the metabolome of each treatment condition are shown. BHB-CoA was not detected except in standards and the MS chromatogram is shown in (G). Data are represented as mean +/- SD and statistical differences were calculated by 1-way ANOVA followed by Tukey's test for multiple comparisons. ****$p<0.0001$. **(H-K)** HEK293T cells were transfected with siRNA and then treated with 10 mM BHB for 4 hours to assess the role of indicated target proteins in Kbhb formation. Kbhb was evaluated by western blot and one representative western blot from N=2 independent experiments is shown in (H). Relative mRNA expression for ACSS1 (I), ACSS2 (J) and ACSS3 (K), which were determined by qPCR analysis. RPL13A was used for normalization. Data are represented as mean +/- SEM of N=2 independently performed experiments. No stastical analysis was applied. **(L)** Western blot of Kbhb abundance in HEK293T cells treated with BHB (10 mM) for 24 hours in the presence of long fatty acyl-CoA synthetase Inhibitor, Triacsin C. Results are representative of N=2 independent experiments.

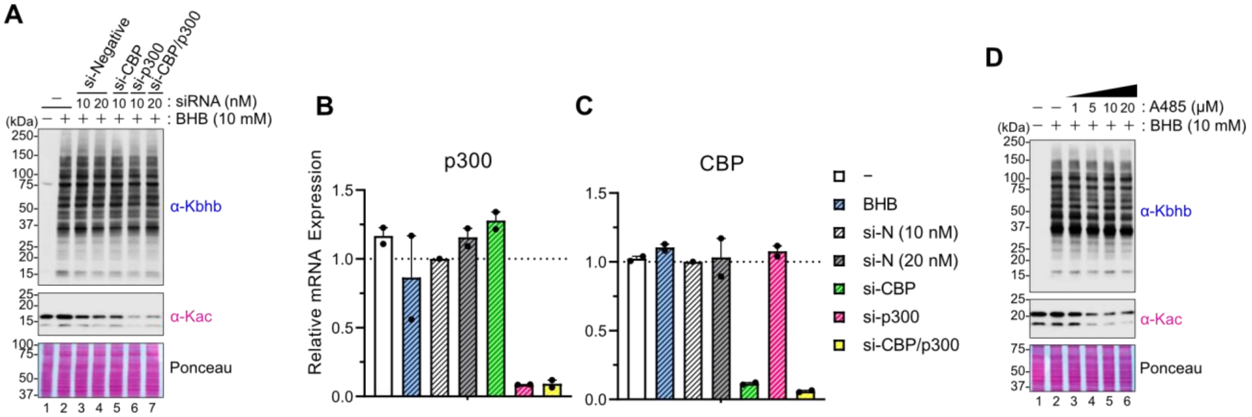


**Figure S4. Global Kbhb formation does not require CBP/p300 acyltransferase activity**

**(A-C)** HEK293T Cells were transfected with siRNA and then treated with 10 mM BHB for 4 hours. Kbhb and Kac abundance were assessed by western blot (A). Relative mRNA expression for p300 (B) and CBP (C) were determined by qPCR analysis. RPL13A was used for normalization. Data are represented as mean +/- SEM of N=2 independently performed experiments. No stastical analysis was applied. **(D)** Kbhb and Kac abundance was measured by western blot of HEK293T cells treated with BHB (10 mM) for 24 hours in the presence of CBP/p300 inhibitor, A485. Data are representative of N=2 independent experiments.

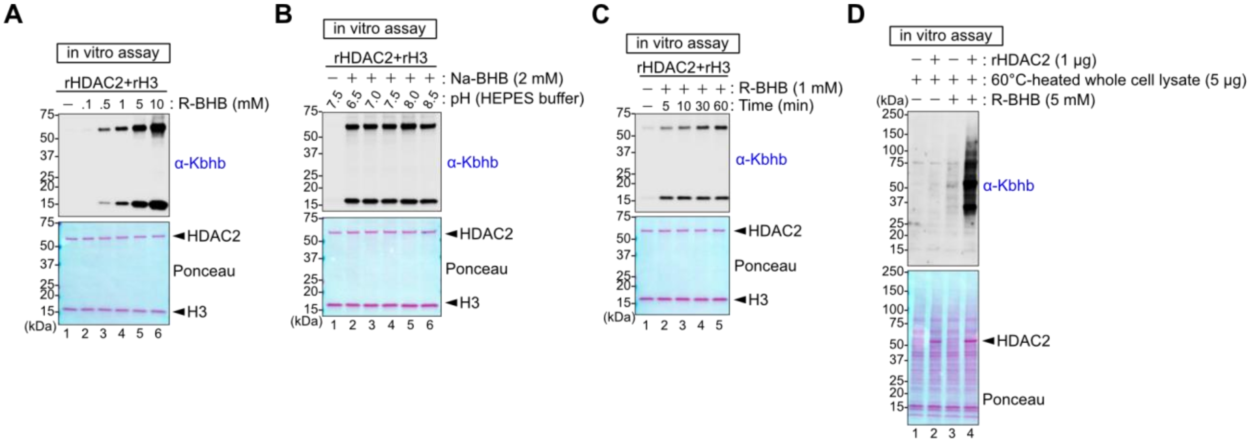


**Figure S5. Characterization of Kbhb dynamics in reconstitution assay**

**(A-C)** Kbhb formation was measured by western blot after in vitro lysine β-hydroxybutyrylation assay with 1 µg rHDAC2 and 1 µg rH3 in the presence of various concentration of R-BHB or Na-D/L-BHB. Reaction was performed at 37°C for 1 hour except for (C). Protein loading was visualized by ponceau S staining. Lysine β-hydroxybutyrylation was detected by anti-Kbhb antibody. HEPES-KOH buffer at the indicated pH was used for the reaction in (B). Reaction was performed at 37°C for the indicated time, from 5 mins to 60 mins, in (C). **(D)** Kbhb formation was measured by western blot in heat-deactivated in whole cell lystates after in vitro lysine β-hydroxybutyrylation assay with 1 µg rHDAC2 in the presence of 5 mM R-BHB. Reaction was performed at 37°C for 1 hour.

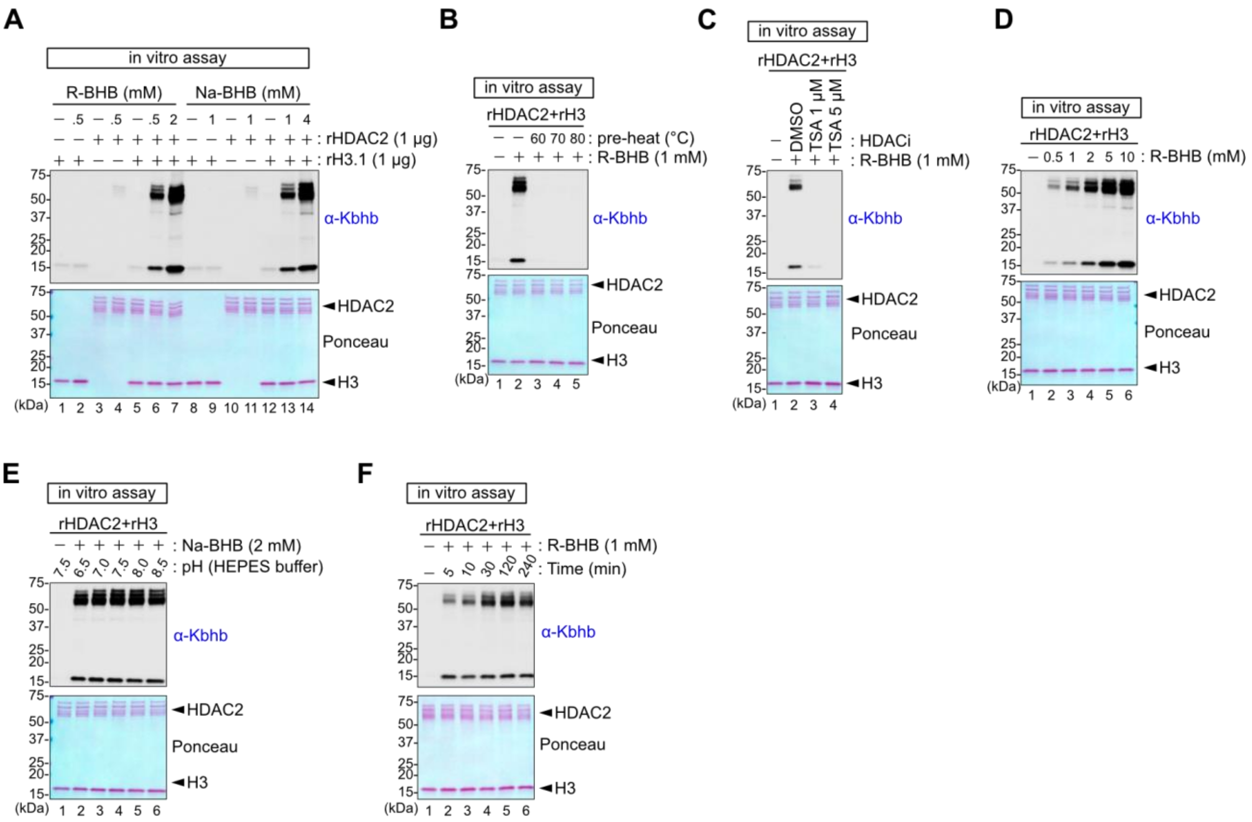


**Figure S6. Lysine β-hydroxybutyrylation activity is observed in alternative commercial source of rHDAC2**

**(A-F)** Repeated experiments of an in vitro lysine β-hydroxybutyrylation assay using rHDAC2 from Active Motif to confirm the reproducibility of the results. Western blots were used to evaluate Kbhb formation of an in vitro lysine β-hydroxybutyrylation assay with recombinant HDAC2 (rHDAC2) and histone H3 (rH3) and R-BHB or Na-D/L-BHB. Reaction was performed at 37°C for 1 hour except for (F). Protein loading was visualized by ponceau S staining. Kbhb was detected by anti-Kbhb antibody. rHDAC2 was inactivated by heating for 1 hour at the indicated temperatures in (B). TSA at 1 or 5 μM was added in the reaction buffer in (C). HEPES-KOH buffer at the indicated pH was used for the reaction in (E). Reaction was performed at 37°C for the indicated time, from 5 mins to 240 mins, in (F).

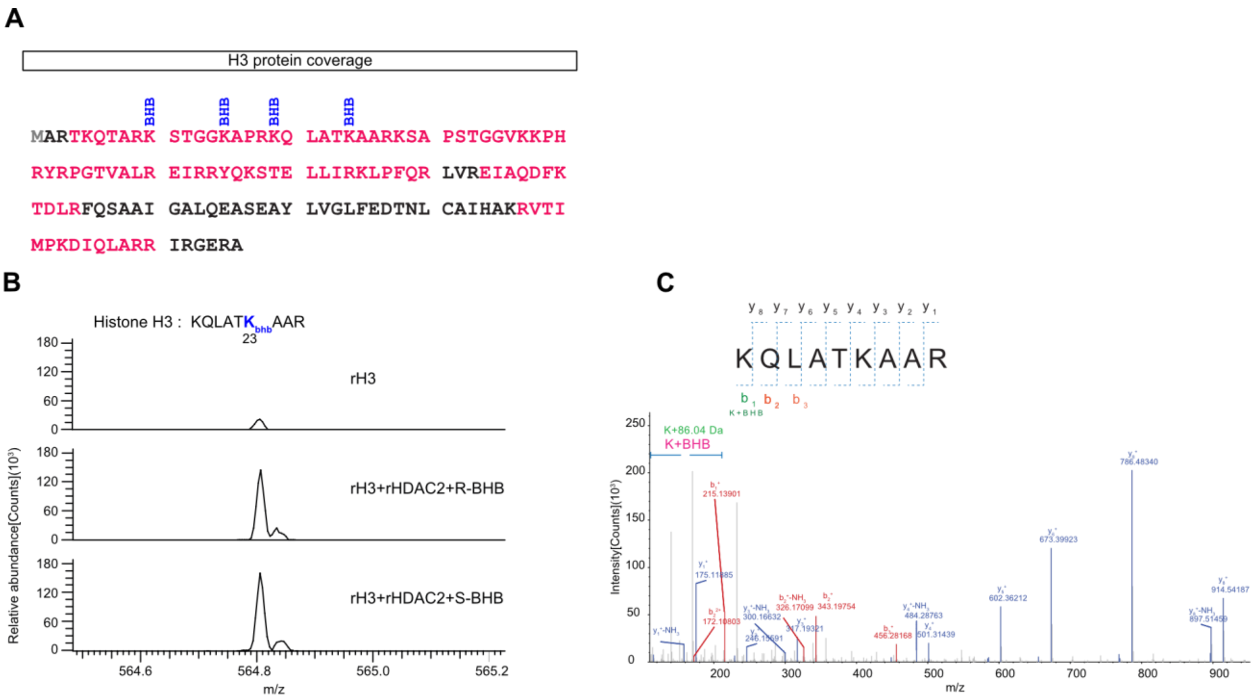


**Figure S7. LC-MS/MS analysis of histone Kbhb peptides after in vitro lysine β-hydroxybutyrylation**

**(A)** The sequence of histone H3.1. We recovered peptides containing all residues highlighted in red text. Kbhb modified peptides are marked by "BHB". **(B-C)** MS-spectrum and MSMS-spectra of the indicated peptide are shown in (B) and (C), respectively.

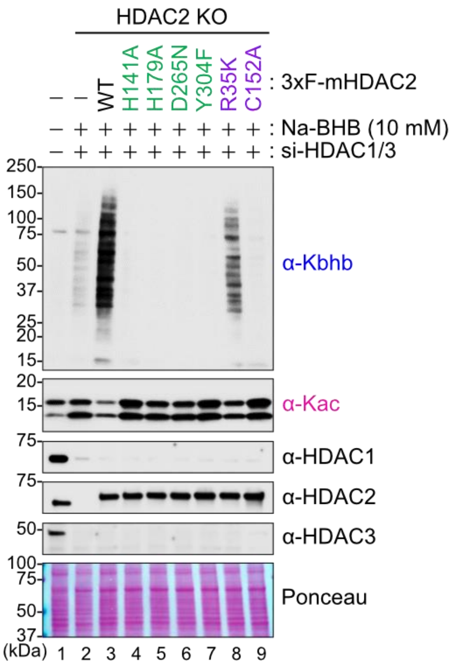


**Figure S8. Establishing stable mutant HDAC2 cell lines**

HDAC2-KO cell lines were complemented with wildtype or indicated 3xFLAG-mHDAC2 mutant proteins. HDAC1 and HDAC3 expression was knocked down by siRNA transfection and cells were treated with 10 mM BHB for 4 hours as indicated. Kbhb, Kac, and HDAC proteins were evaluated by western blot. Blots come from a single experiment and are representative of N=3 independent experiments.

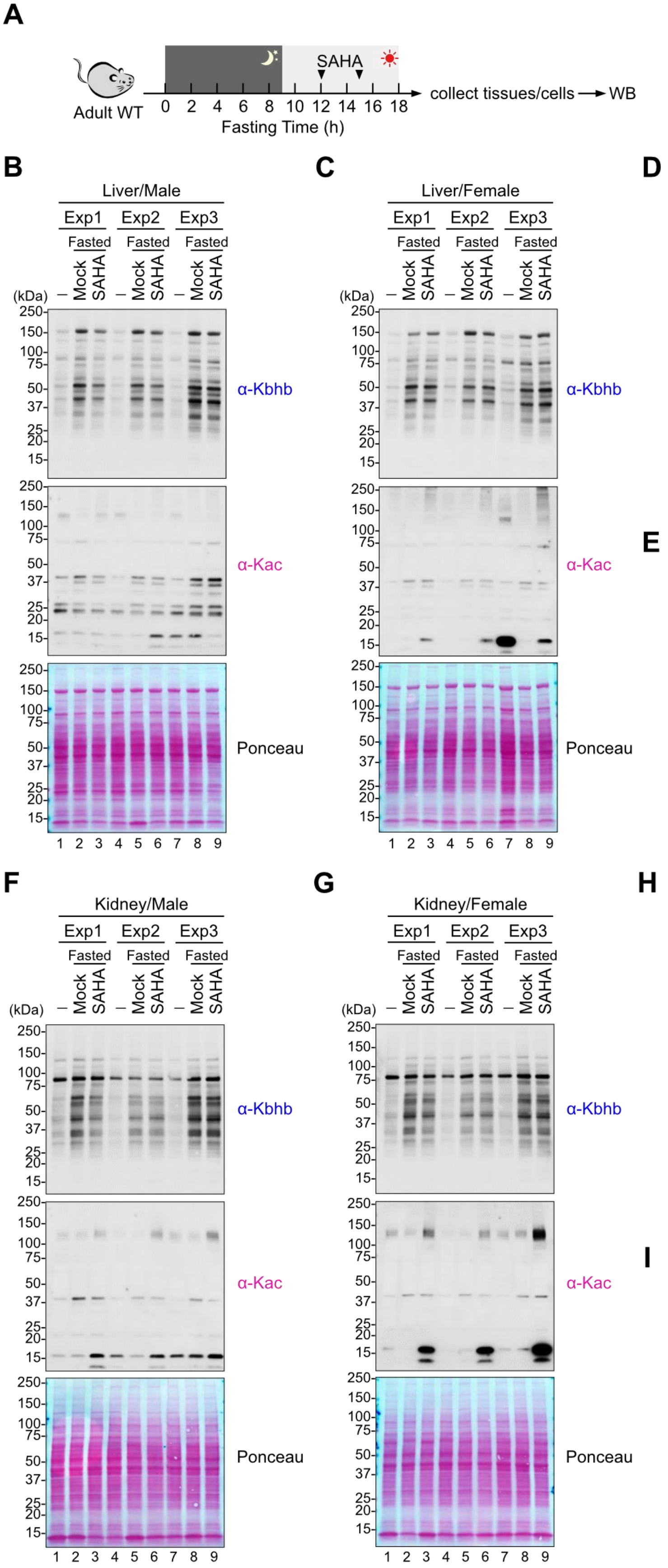

A
Adult WT
SAHA
0 2 4 6 8 10 12 14 16 18
Fasting Time (h)
collect tissues/cells → WB
B
Liver/Male
Exp1 Exp2 Exp3
Fasted Fasted Fasted
– Mock SAHA – Mock SAHA – Mock SAHA
(kDa)
α-Kbhb
α-Kac
Ponceau
1 2 3 4 5 6 7 8 9
C
Liver/Female
D
Kbhb
(Liver)
Relative Intensity
Mock SAHA
E
Kac
(Liver)
*
F
Kidney/Male
G
Kidney/Female
H
Kbhb
(Kidney)
I
Kac
(Kidney)

**Figure S9. Kbhb induction in the liver and kidney from SAHA-injected mice**

**(A)** The schematic of experimental design is same as Fig. 5A. C57BL/6 mice were fasted starting from night time and injected SAHA twice at 12 and 15 hr after fasting was initiated. WB: wetern blot analysis. **(B-E)** Western blots of liver are shown in (B-C). Exp1-3: N=3 independent experiments are shown on each blot, each containing a fed control, a mock-treated, and a SAHA-treated mouse as indicated. The signals were normalized by ponceau S staining and fold-changes relative to Fed sample for Kbhb and Kac are plotted in (D) and (E), respectively. Circles indicate males and triangles indicate females. Unpaired t-test with two-tailed P values and lines connect the control and SAHA-treated mice from the same independent experiment. **(F-I)** Western blots of kidney are shown in (F-G). Exp1-3: N=3 independent experiments are shown on each blot. The signals were normalized by ponceau S staining and fold-changes relative to Fed sample for Kbhb and Kac are plotted in (H) and (I), respectively. Circles indicate males and triangles indicate females. Unpaired t-test with two-tailed P values and lines connect the control and SAHA-treated mice from the same independent experiment. *$p<0.05$
